# Pinned, locked, pushed, and pulled traveling waves in structured environments

**DOI:** 10.1101/341222

**Authors:** Ching-Hao Wang, Sakib Matin, Ashish B. George, Kirill S. Korolev

## Abstract

Traveling fronts describe the transition between two alternative states in a great number of physical and biological systems. Examples include the spread of beneficial mutations, chemical reactions, and the invasions by foreign species. In homogeneous environments, the alternative states are separated by a smooth front moving at a constant velocity. This simple picture can break down in structured environments such as tissues, patchy landscapes, and microfluidic devices. Habitat fragmentation can pin the front at a particular location or lock invasion velocities into specific values. Locked velocities are not sensitive to moderate changes in dispersal or growth and are determined by the spatial and temporal periodicity of the environment. The synchronization with the environment results in discontinuous fronts that propagate as periodic pulses. We characterize the transition from continuous to locked invasions and show that it is controlled by positive density-dependence in dispersal or growth. We also demonstrate that velocity locking is robust to demographic and environmental fluctuations and examine stochastic dynamics and evolution in locked invasions.

## Introduction

Propagation of waves, fronts, and pulses is a recurrent theme in natural sciences [1]. In evolution, they describe the geographic spread of a beneficial allele or the increase of fitness over time [2–4]. In ecology, they capture epidemics, invasions of foreign species, and range shifts due to a climate change [5–7]. In cell biology, traveling excitations describe electrical pulses in neurons, calcium waves in various tissues, and the assembly of large cytoskeletal complexes [1, 8–13]. The list of applications also includes crystal growth, combustion fronts, and even the spread of entanglement in quantum mechanics [14–16].

Given the diverse settings in which traveling fronts occur, numerous approaches have been developed to model front propagation in specific systems. Most of these approaches can be grouped into four classes depending on whether space and time are treated as discrete or continuous (Fig. 1). Discrete time typically represents non-overlapping generations or periodic seasonal forcing, while discrete space accounts for habitat fragmentation.

**Figure 1.**
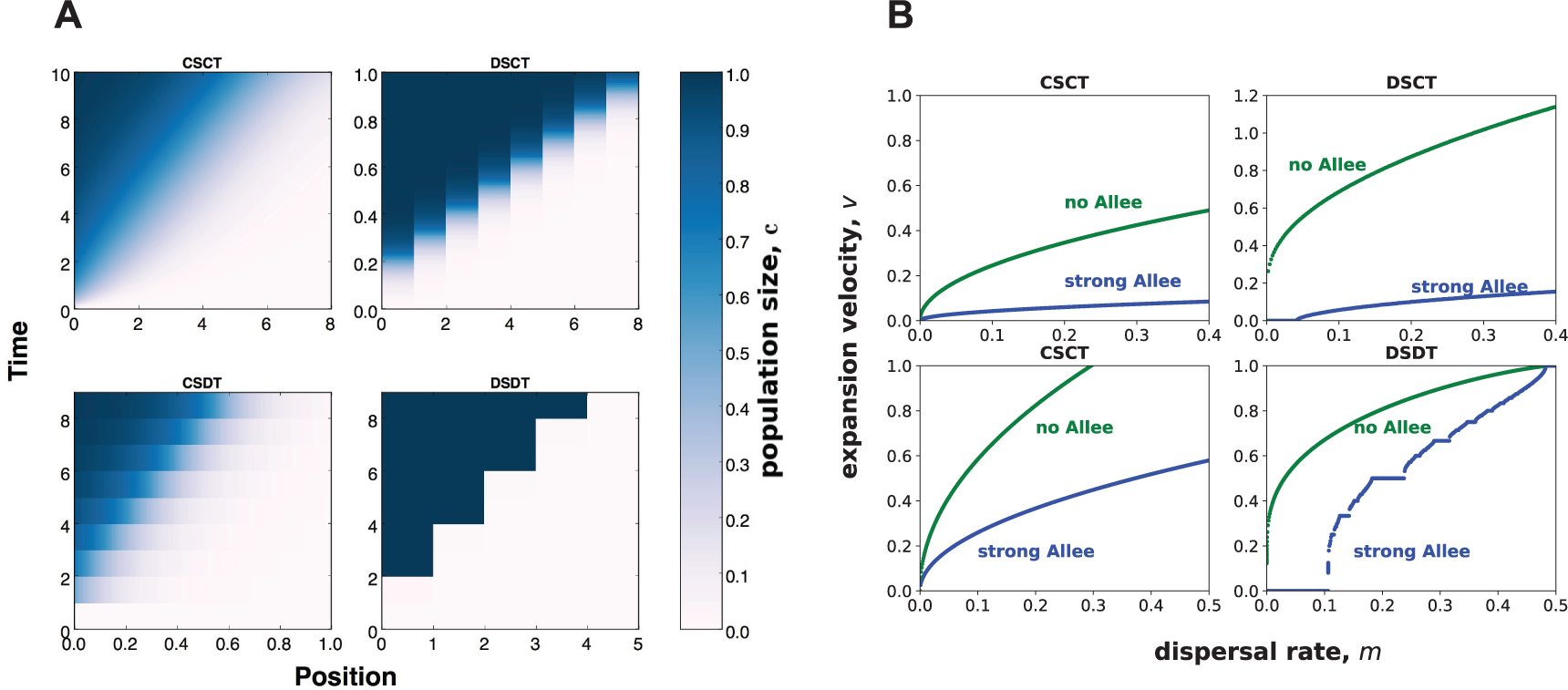
Discrete and continuous models predict distinct invasion dynamics. Both panels compare four types of models that differ in whether space (S) or time (T) are treated as discrete (D) or continuous (C). **(A)** Species (blue) expands into empty space (white) starting from a step-like initial condition. Sharp changes in population density indicate discreteness in the underlying model. **(B)** The dependence of the expansion velocity *v* on the migration rate *m* for the same four models. When the space is continuous *v*(*m*) is a smooth function. The same behavior is observed in models with fragmented landscapes but continuous-time growth (DSCT), except a critical migration rate might be necessary for the invasion to proceed. In contrast, the dependence of *v* on *m* could be highly non-analytic in models with discrete space and time (DSDT), which manifests in constant velocities over a large range of migration rates. Note that all four models exhibit smooth *v*(*m*) in the absence of positive density-dependence (Allee effect). Even for smooth *v*(*m*), there is a difference between the continuous-space models, for which *v*(*m*) ~ *m*^1/2^ for small *m*, and discrete-space models, for which *v*(*m*) is more singular at the origin [17]. Parameter values for the four models are provided in Methods.

Continuous time, continuous space (CTCS) models arise naturally for chemical reactions and are typically formulated in terms of partial differential equations. In ecology and evolution, CTCS models describe population with overlapping generations living in a spatially homogeneous habitat. The simplest mathematical formulation of a CTCS model reads
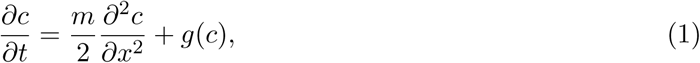

where *c*(*t, x*) is the population density that depends position *x* and time *t; m* is the migration or dispersal rate; and *g*(*c*) is the growth rate of the population.

Continuous space, discrete time (CSDT) models describe species with strong seasonality such as annual plants that spread in a spatially homogeneous landscape. These models are expressed in terms of integro-difference equations that specify how population densities change from generation to generation:
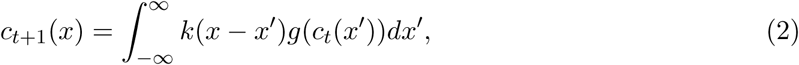

where *c_t_*(*x*) is the population density as a function of position *x* at discrete generation *t; k*(*x* − *x*′) is the dispersal kernel that specifies the probability of dispersal to position *x* from position *x*′; and *g*(*c_t_*(*x*)) describes the growth dynamics. The kernels that are sufficiently short-range, e.g. exponential or Gaussian, result in an invasion front that propagates at a constant speed. Fat-tailed kernels, e.g. with power law tails, could result in accelerating invasions [18–20].

The assumption of continuous space does not hold for many ecological populations. Patchy habitats arise due to a low density of locations suitable for growth, due to turbulent flows in marine environments [21, 22], and due to internal ecological dynamics that create spatial patterns via Turing instabilities, ecological drift, and other mechanisms [23–26]. Human development also leads to habitat fragmentation [27] with row and grid planting of agricultural plants being extreme example of regular spatial structures. In addition, habitats with regular arrangement of patches are often engineered using microfluidics or other technologies to create microcosm metapopulations for experimental study of collective behavior, ecology, and evolution using laboratory populations [28–33]. Outside ecology, spatial structures arise due to crystal lattices, clustering of ion channels, or spatially periodic flow patterns in reaction chambers [12, 14, 16, 34].

Discrete space, continuous time (DSCT) models are appropriate for fragmented landscapes with little or no seasonal forcing. The dynamics are described by a set of ordinary differential equations coupled by dispersal, which, in the simplest case, occurs only between the nearest patches:
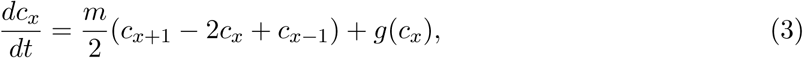

Finally, seasonal growth in fragmented landscapes is described by discrete space, discrete time (DSDT) models expressed as difference equations, which are also known as coupled map lattices and cellular automata:
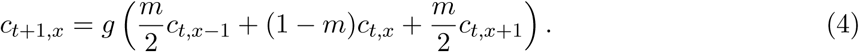

In addition to natural populations, DSDT models capture the dynamics in some experimental studies [28, 29] and underlie many simulations because discretization is often used in numerical methods.

Despite significant differences, the models in all four classes can describe the propagation of a traveling front. As a result, the choice of the model is often dictated not only by its relevance to a specific population, but also by its mathematical or computational tractability [35–37]. Reaction-diffusion equations are commonly used to obtain analytical insights into population dynamics [1, 38–43], and integro-difference equations are preferred when it is necessary to account for long-range dispersal [18, 19]. Models with discrete time or space also have certain advantages; for example, they provide a convenient way to introduce stochastic dynamics or use satellite data on natural landscapes [37, 44, 45].

Some phenomena, however, arise most naturally only in a specific model class. One important example in invasion pinning (Fig. 1), which occurs when the front gets stuck at a particular patch or landscape heterogeneity [17, 46]. Invasion pinning requires habitat fragmentation and the presence of a strong Allee effect, i.e. a threshold density below which the growth rate is negative. Under these conditions, a critical dispersal rate is required to initiate population growth in neighboring patches. Above the critical dispersal rate, invasion proceeds as in continuous-space models, but, below it, invasion front is pinned.

In models where both time and space are discrete, invasions can become not only pinned, but also locked. Locked fronts advance periodically in a pulsed fashion, and their velocities assume only a discrete set of values, which are insensitive to small changes in dispersal and growth rates (Fig. 1).

Although locking of traveling fronts may seem peculiar, it has been observed for the Belousov-Zhabotinsky reaction spreading in a periodic array of periodically-driven vortices [34]. Moreover, the characteristic pattern of plateaus shown in Fig. 1 was found in a variety of physical systems including arrays of Josephson junctions [47], chemical reactions [48], quantum gases [49], and crystal structures of alloys [50]. This pattern, dubbed a Devil’s staircase [51], is a manifestation of a phenomenon known as mode locking in nonlinear sciences. In the simplest case, mode locking occurs when the observed frequency of a nonlinear oscillator equals a rational number times the frequency of the external drive. The plateaus are regions of the drive amplitude that correspond to the same ratio of the oscillator and drive frequencies.

The theory of mode locking has been applied to coupled map lattices to explain the locking of invasion velocities, and some rigorous results are available on the existence and properties of locked fronts [52–58]. Previous studies, however, lacked ecological context and focused exclusively on bistable maps. Therefore, they did not explore the transition from continuous to locked fronts as the strength of an Allee effect or other ecological parameters are varied (Fig. 1). Understanding this transition and the differences between DSDT and other models are the primary goals of this paper.

We show that velocity locking requires positive density-dependence in growth or dispersal, but bistability is not necessary. In addition to locked and pinned waves, we report two types of continuous expansions in DSDT models. These types are analogous to pulled and pushed waves in reaction-diffusion equation [29, 39, 59]. Our results further demonstrate that velocity locking occurs both in one and two spatial dimensions and is robust to significant levels of demographic and environmental fluctuations. The effects of velocity locking on stochastic front wandering and evolutionary dynamics are also discussed.

## Results

### Velocity locking occurs in many ecological models

We studied a variety of ecological growth models to determine whether velocity locking is a generic phenomenon. One convenient choice of *g*(*c*) is the piecewise-linear growth model:
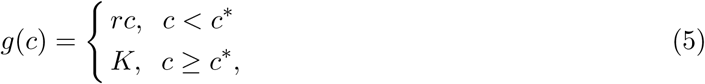

where *r* is the growth rate at low densities, *K* is the carrying capacity, and *c*\* is the density sufficient to reach the carrying capacity. When *rc*\* = *K*, the growth function is continuous, and there is no Allee effect because the per capita growth rate *g*(*c*)/*c* monotonically decreases with population density. When *rc*\* < *K*, there is a jump in *g*(*c*) at *c* = *c*.* This jump corresponds to an increase in the growth rate due intraspecific facilitation, which requires a certain population density [60, 61].

Because of the jump, *g*(*c*)/*c* is non-monotonic, and the growth dynamics are said to exhibit an Allee effect [60, 61]. When *r* > 1, the Allee effect is termed weak. In this case, small populations always grow. When *r* < 1, the Allee effect is said to be strong, and small populations go extinct unless they are rescued by immigration from nearby patches. In the following, we often use this piecewise-linear growth model because it is mathematically tractable and allows one to easily tune the strength of the Allee effect.

Although Eq. (5) is a reasonable model of ecological dynamics, it not clear whether velocity plateaus in this model arise due to discontinuities in *g*(*c*). To answer this question and determine whether velocity locking is a generic phenomenon that is not specific to a particular *g*(*c*), we considered two alternative growth functions that had complementary properties and covered a wide range of ecological scenarios. Note, however, that we only considered growth functions that describe a steady approach to the carrying capacity and do not lead to chaotic or periodic dynamics on their own.

The Beverton-Holt model with offset [62, 63] is commonly used to model population dynamics, e.g. of fisheries, and is specified by the following equation:
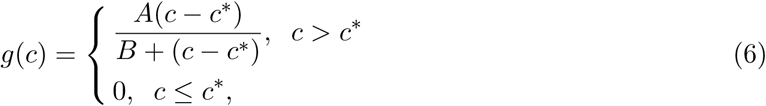

where *A* sets the carrying capacity, *B* sets the density at which intraspecific competition starts to significantly affect population growth, and *c*\* sets the magnitude of the Allee effect.

Although *g*(*c*) in the Beverton-Holt model is continuous, it has a discontinuous derivative at *c*.* To exclude the possibility that this non-analyticity is responsible for velocity locking, we considered an infinitely differential map, known as Hill function in molecular biology [64]. Hill-like functions have also been used to model population dynamics in ecology [65]. The simplest Hill function reads
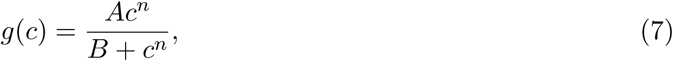

where *A* and *B* play the same role as in the Beverton-Holt model, and *n* controls the strength of the Allee effect. The Allee effect is absent for *n* = 1 and increases with *n* for *n* > 1.

All three models exhibited velocity locking and showed clear plateaus in the dependence of *v* on *m* (Fig. 2). Thus, velocity locking is not caused by singularities in the growth function, although velocity plateaus are larger for discontinuous or rapidly varying *g*(*c*). We then conclude that velocity locking is robust to the choice of modelling assumptions, control parameters, and differences in species ecology. Therefore, it should be expected for a generic growth model.

**Figure 2.**
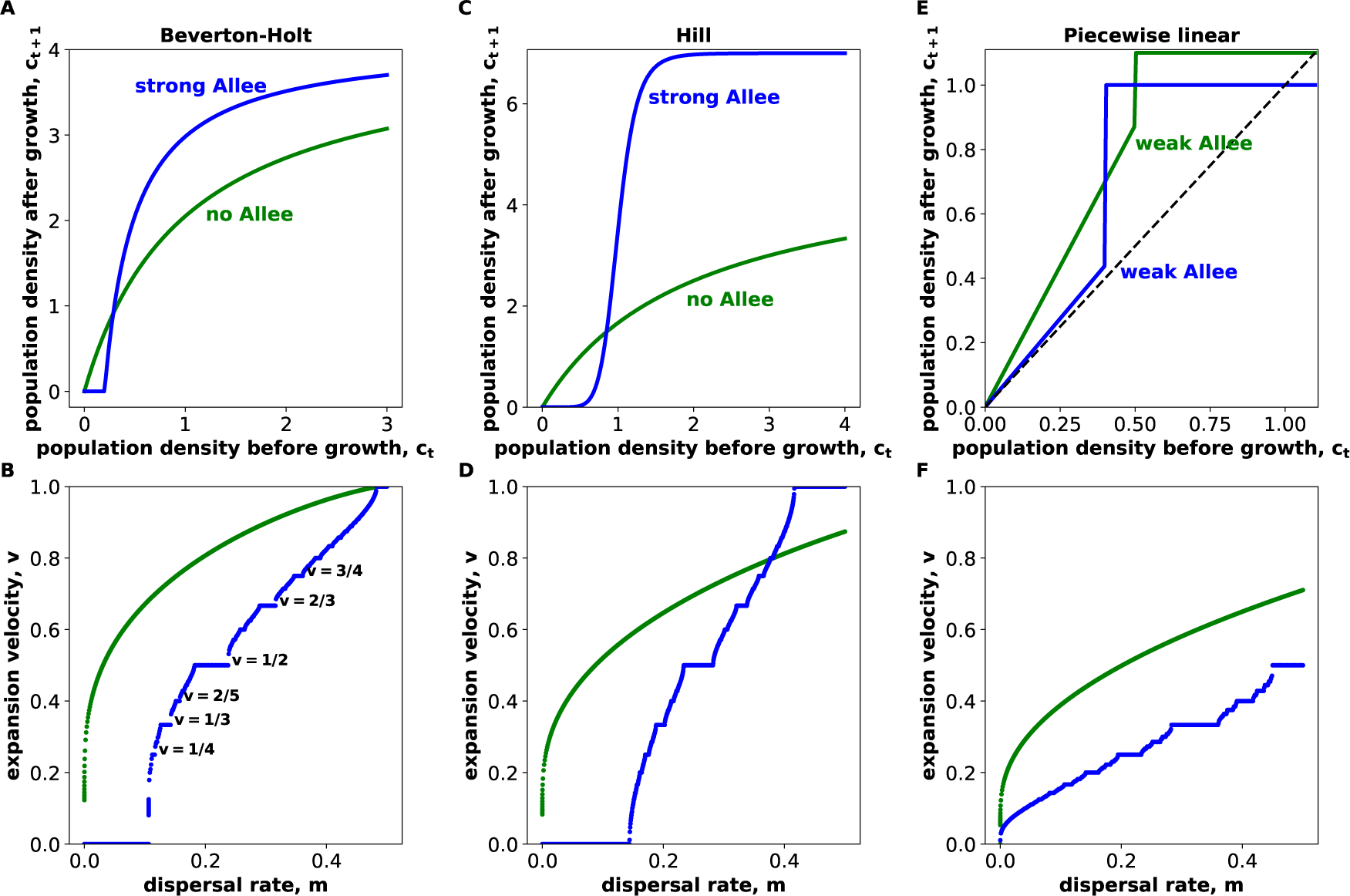
Velocity locking occurs in many ecological models. The growth functions *g*(*c*) for three ecological models are shown in the top three panels **(A, C, E).** Below each model, we plotted the dependence of the invasion velocity on the dispersal rate **(B, D, F).** Panels B and D illustrate the fact that velocity locking occurs in the presence of a strong Allee effect, but not when an Allee effect is absent. Panels E and F show that velocity locking can occur for a weak Allee effect, but only if it is sufficiently large. For BevertonHolt model, we used *A* = 4.1, *B* = 0.3, and *c*\* = 0.2 for the strong Allee effect conditions and *A* = 4.1, *B* = 1, and *c*\* = 0 for the no Allee effect conditions. For the Hill model, we used *A* = 7, *B* = 1, and *n* = 8 for the strong Allee effect conditions and *A* = 5, *B* = 2, and *n* = 1 for the no Allee effect conditions. For the piecewise-linear model, we used *r* = 1.1, *K* = 1, and *c*\* = 0.5 for the weak Allee effect conditions and *r* = 1.1, *K* = 1.0, and *c*\* = 0.4 for the weak Allee effect conditions.

### Locked invasions are periodic

Velocity locking invariably results in a periodic and often pulsed propagation of the invasion front (Fig. 3). The pulsations occur because several generations of slow growth and migration are necessary to reach the density at which intraspecific facilitation enables rapid growth. After this rapid growth, the density profile assumes exactly the same shape as in the beginning of the cycle. In general, locked fronts advance by *p* patches every *q* generations [52, 53], and there are only *q* distinct density profiles: one for each generation from *t* to *t*+*q* (Fig. 4). As a result, invasion velocities take only rational values, *v* = *p/q*, when measured in units of inter-patch distance per generation (Fig. 2). The periodicity of front motion remains invariant within the velocity plateau even though the shape of the front does change with the model parameters.

**Figure 3.**
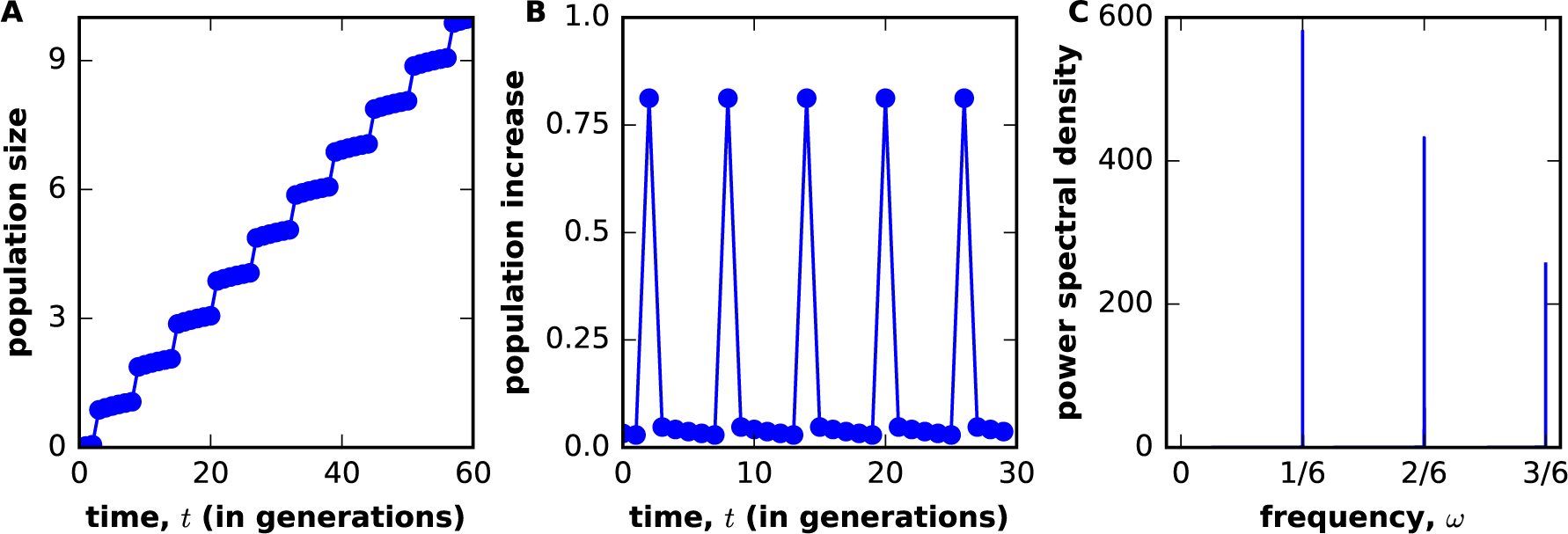
Locked fronts advance by periodic pulses. **(A)** shows how the total population size of the invader increases with time for *v* = 1/6 velocity plateau in the piecewise-linear growth model. The staircase-like increase of the population size indicates that the invasion is pulsed. This is more clearly visible in **(B),** where the change in the population size per generation is plotted. The pattern is clearly periodic, and most of the growth occurs during only of the one generations in a period. Thus, the invader spreads in pulses that repeat every six generations. **(C)** confirms the periodicity of the invasions by showing the power spectrum of the population size change relative to its mean over the period. The peaks at the fundamental frequency of 1/6 and higher harmonics are clearly visible. Here, *m* = 0.110, *r* = 0.93, and *c*\* = 0.22.

**Figure 4.**
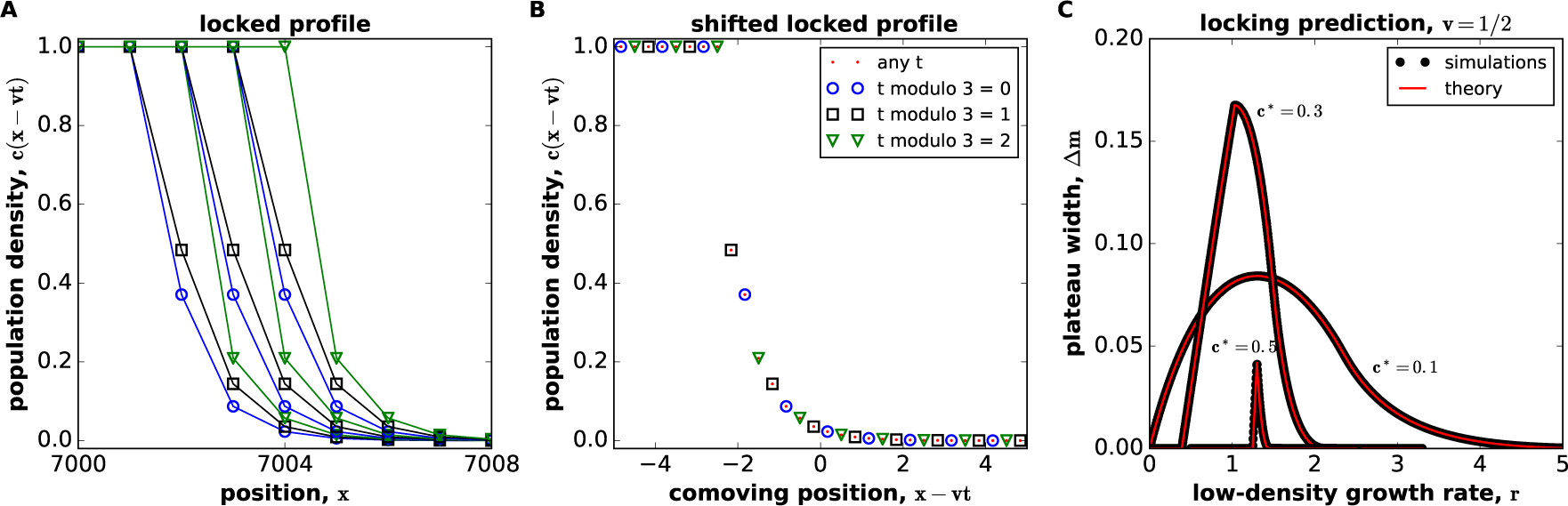
Locked invasions have discrete profiles that repeat periodically. **(A)** shows the population density profile at nine consecutive generations for *v* = 1/3 plateau in the piecewise-linear model. To highlight the fact that density profiles repeat periodically, we used different symbols for different generations modulo 3. **(B)** Density profiles collected over 10^3^ generations were shifted by −*vt* to transform them into the reference frame co moving with the invasion. The resulting profile is discrete with only a countable number of distinct values of *c.* **(C)** The width of *v* = 1/2 plateau is shown as a function of *c*.* There is an excellent agreement between simulations and the theoretical calculation that assumes that front shape repeats periodically (see Methods). This agreement further confirms the periodic nature of locked invasions. Note that a single plateau can occupy a very large part of the range of possible dispersal rates. In A and B, *r* = 1.1, *K* = 1, *c*\* = 0.5, and *m* = 0.4. In C, *K* = 1 and other parameters are varied.

To verify the periodic nature of locked waves, we obtained an exact solution for the invasion dynamics inside *v* = 1/2 plateau for the piecewise-linear growth model (see Methods). Our solution agrees both with the population density profiles and with the locations of the velocity plateaus obtained in simulation (Fig. 4C). This agreement demonstrates that Eq. (4) admits a moving front with the same periodicity over a region of model parameters and thus firmly establishes that *v* is exactly rather than approximately constant within the plateau. More importantly, our calculation confirms that velocity locking is a result of synchronization of the invasion dynamics with the spatio-temporal periodicity of the habitat.

### Pulled waves propagate without locking in DSDT models

Not all invasions in DSDT models are locked; see Figs. 1 and 2. In fact, it can be rigorously demonstrated that *v*(*m*) is a smooth function for so-called pulled waves, which are driven by the growth and dispersal at the very tip of the invasion front. Invasions are guaranteed to be pulled when there is no Allee effect, and, therefore, the per capita growth rate is maximal at the leading edge of the front [3, 37, 59, 66]. Because *c* is small at the front, nonlinear terms in *g*(*c*) are negligible, and the invasion velocity can be computed exactly by linearizing Eq. (4) and then solving it via Laplace and Fourier transforms [59]; see Methods.

The main result of this calculation is that pulled waves in DSDT models behave as in CSCT models. Specifically, the population density at long times depends only on *x* − *vt* and is described by a continuous function *c*(*x* − *vt*) with *v* given by
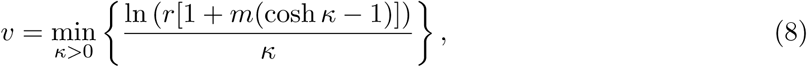

where *r* = lim_c→0_ *g*(*c*)/*c.*

In the absence of an Allee effect, numerical simulations show excellent agreement with the theory of pulled waves. The invasion velocities predicted by Eq. (8) perfectly matched the simulation results for the piecewise-linear, Beverton-Holt, and Hill model (Fig. 5A). In addition, the observed population densities formed a continuous profile after a shift by *vt* to transform them into the reference frame comoving with the invasion (Fig. 5B). Note that, for locked waves, the shift to the comoving reference frame produces a discrete profile (Fig. 4B) because there are only *q* distinct profiles. We therefore conclude that pulled waves are not locked, and DSDT models support both periodic and aperiodic invasions.

**Figure 5.**
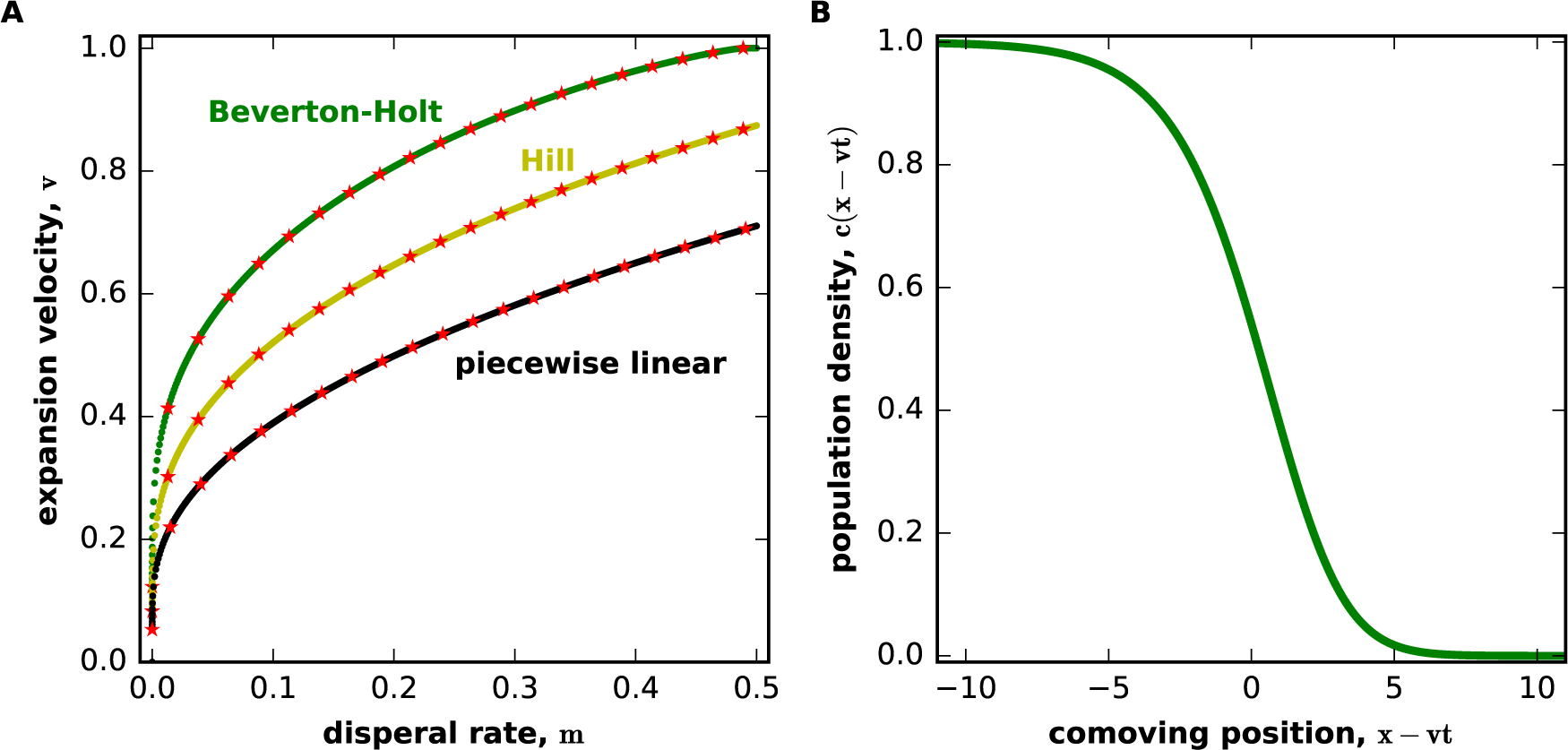
Pulled waves propagate without velocity locking. **(A)** The dependence of the invasion velocity on the dispersal rate is shown for Beverton-Holt, Hill, and piecewise-linear growth models from Fig. 2. The theoretical prediction for pulled waves from Eq. (8) is shown with red stars. The excellent agreement between the theory and simulations confirms that models species can spread without velocity locking in DSDT. **(B)** is similar to Fig. 4B and shows that population density profiles obtained at different times collapse onto a continuous curve after a shift by −*vt* to transform them into a comoving reference frame. This collapse indicates that, in steady state, *c*(*t, x*) = *c*(*x* − *vt*) for pulled waves. For this panel, the Beverton-Holt growth map was used with *A* = 3, *B* = 2, *c*\* = 0, and *m* = 0.5.

### Pushed waves: A second type of unlocked expansions in DSDT models

To understand the transition from locked to unlocked fronts, we examined how the invasion dynamics change with the strength of an Allee effect. For reaction-diffusion waves in CSCT models, a critical strength of an Allee effect is necessary to transform pulled waves into a new spreading regime [29, 39, 59]. Invasions in this regime are known as pushed waves because nonlinearities in *g*(*c*) contribute to front propagation and, therefore, the spreading velocities are greater than predicted by Eq. (8). We found similar behavior in DSDT models. Figure 6 shows that Eq. (8) remains valid even in the presence of a substantial Allee effect, but the observed velocities become greater than the prediction of the linear theory once a critical strength of the Allee effect is reached.

**Figure 6.**
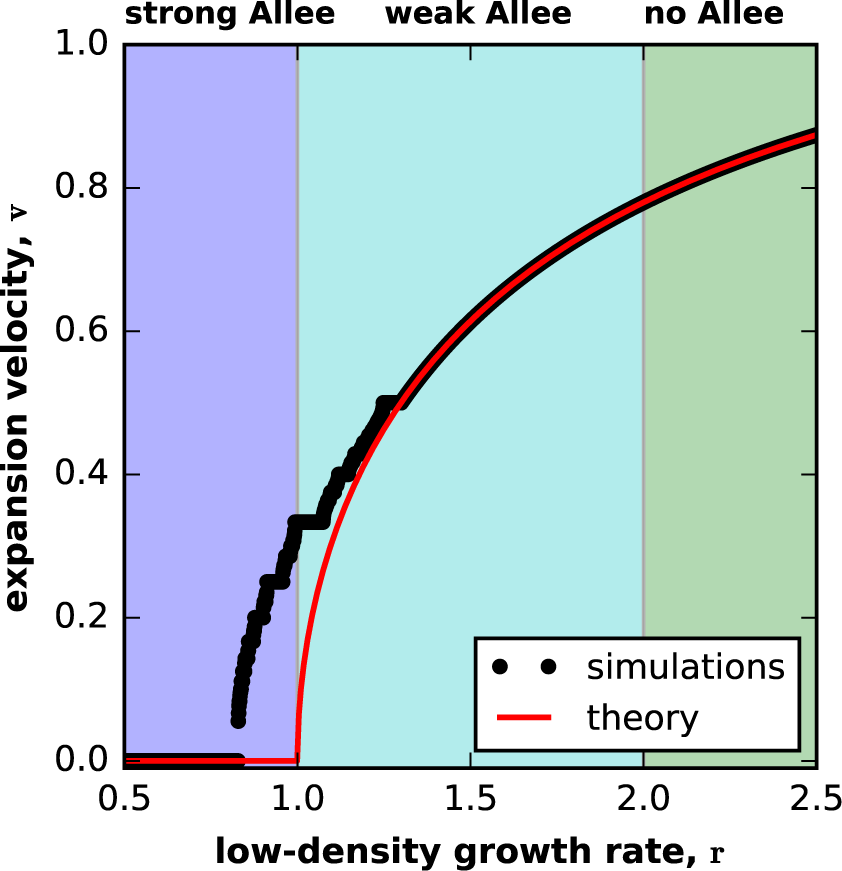
Expansions stop being pulled at intermediate strengths of a weak Allee effect. Black dots show the dependence of the invasion velocity on the low density growth rate for the piecewise-linear model, and the red line shows the prediction of Eq. (8). For high *r*, the data and theory overlap indicating that invasions are pulled. For smaller *r*, there is disagreement and clear signs of velocity locking. The transition between the two regimes occurs at *r* ≈ 1.3, which is substantially lower than the critical growth rate that marks the onset of a weak Allee effect (*r* = 2.0). Here, *K* = 1, *c*\* = 0.5, and *m* = 0.5.

We then asked whether locked and pulled waves are the only two classes of expansions in DSDT models or whether DSDT models also support unlocked waves that are pushed rather than pulled. To answer this question, we obtained the phase diagram of invasion velocities in the piecewise-linear growth model. Specifically, we computed *v* for (*r, m*) ∊ (0, 4) × (0, 0.4) from simulations with *r* and *m* varied in steps of *δr* = 0.004 and *δm* = 0.0004. The results are shown in Fig. 7A. Waves were labeled as pulled if their velocity differed from Eq. (8) by less than 10^−5^. Locked waves were defined as waves whose velocity differed by less than 10^−5^ from the velocities in at least one of the neighboring simulations, i.e. simulations with parameters changed by *δr* or *δm.* Locked invasions with *v* < 10^−5^ were labeled as pinned. Our main result is that some parameter combinations for which the invasions can not be classified as either pulled or locked. That is their velocities appear to change continuously with model parameters, yet the velocities are greater than the expectation for pulled waves. In analogy with CSCT models, we label such invasions as pushed.

**Figure 7.**
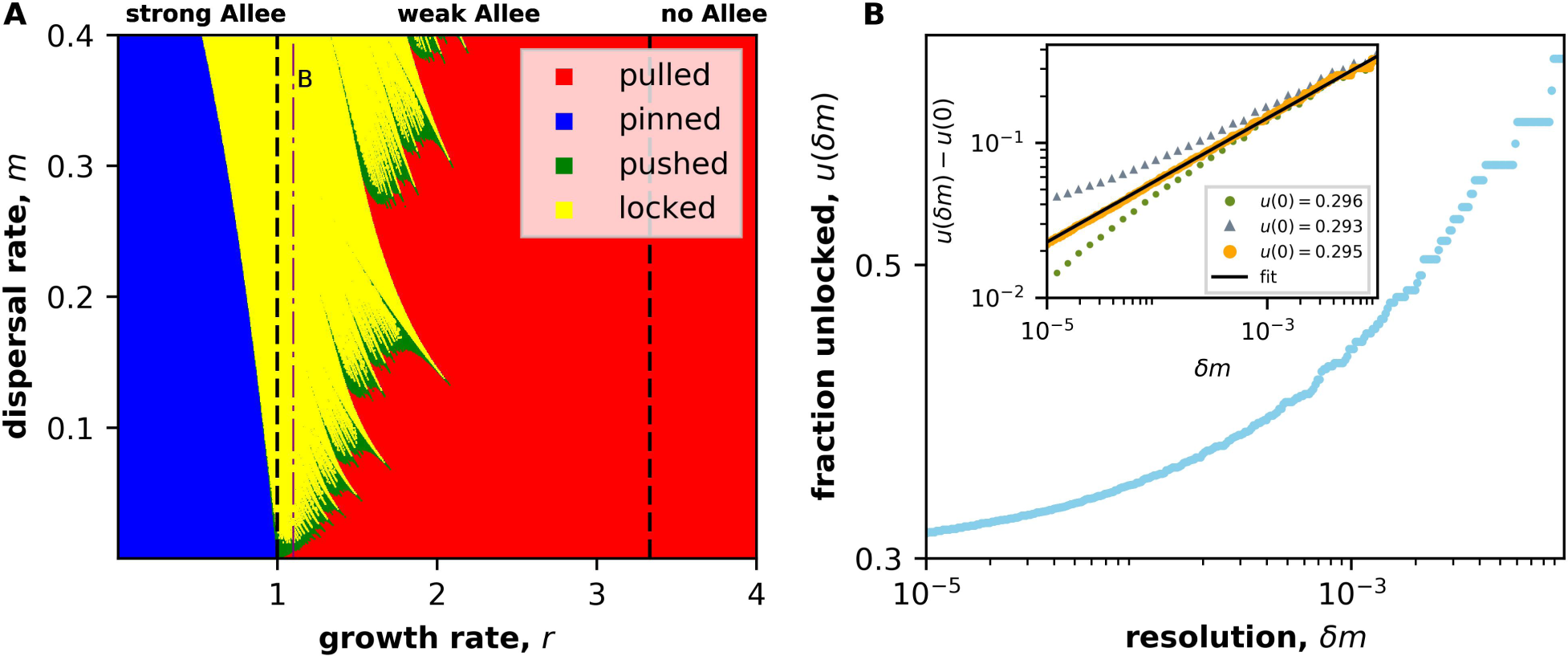
Pinned, locked, pushed, and pulled invasions in DSDT models. **(A)** The phase diagram of invasion classes for the piecewise-linear model with *c*\* = 0.3 and *K* = 1.0. The velocity was computed from simulations where *r* and *m* were varied in steps of *r* = 4. 10^−3^ and *δm* = 4. 10^−4^. Waves were labeled as pulled if their velocities agreed with Eq. (8) with absolute error less than 10^−5^. Waves were classified as locked if their velocities differed by less than 10^−5^ from the velocities obtained in one of the neighboring simulations. **(B)** (Main plot) The fraction of unlocked waves seems to approach a finite limit of infinite resolution *δm* = 0; here *r* = 1.1 corresponding to the thin purple line in A. (Inset) For small *δm*, the fraction of unlocked waves behaves as a power law: *u*(*δm*) ≈ *u*(0) + *δm^ß^.* The data are shown with yellow symbols, and the black line indicates the fit. The best fit is obtained with *u*(0) = 0.295 and *ß* = 0.4, using the python function scipy. optimize. curve fit. Green and grey symbols show that small changes in *u*(0) destroy the clean power law and thus demonstrate that our estimate of *u*(0) is accurate. Note that *u*(0) is significantly greater than the fraction of pulled waves along the purple line in the phase plot, which is equal to 0.014. Hence, some waves must be pushed.

In principle, the existence of pushed waves in Fig. 7A could be attributed to finite *δr* and *δm* in our simulations. Indeed, velocity plateaus smaller than *δr* and *δm* are missed by the procedure described above. To account for this possibility, we considered a slice of the phase diagram at *r* = 1.10 and examined how the fraction of unlocked waves *u* scales with *δm*; see Fig. 7B. Consistent with similar analyses of mode locking in nonlinear oscillators [67–69], we found that *u*(*δm*) ≈ *u*(0) + *δm^β^* with *β* = 0.4 and *u*(0) = 0.295. Importantly, *u*(0) is significantly greater than the fraction of pulled waves, 0.014 (the latter number is much less sensitive to *δm* and, therefore, does not require interpolation to *δm* = 0). This analysis suggests some of the waves at *r* = 1.1 and *m* ∊ (0, 0.4) must be pushed.

The transitions between different modes of propagation in DSDT models is more complex than in other models. For CSCT model, the transition between pulled and pushed waves occurs at a specific strength of the Allee effect and is independent of *m.* In contrast, the boundary between different regimes depends on both *r* and *m* in DSDT models. Moreover, these boundaries have fractal geometry due to an infinite number of Arnold tongues, i.e. regions in parameter space corresponding to the same velocity plateau [67, 70]. Each Arnold tongue has a smooth boundary, including the tongue for *v* = 0, which corresponds to the region of pinned waves.

### Response to perturbations

To better understand the differences between pulled, pushed, and locked invasions, we examined their response to perturbations. In particular, we determined how the effect of a small perturbation decays in time and whether the system returns to unperturbed state in the long time limit. Perturbations were implemented after the invasion reached a steady state by replacing *c* with *c*+*ϵc*(1−*c*). We then examined the difference in *c*(*t, x*) for perturbed and unperturbed copies of the system.

We expected that a perturbation of pulled and pushed fronts should lead to a nonzero shift of the front profile. This expectation follows from the following argument. The movement of unlocked fronts is described by a continuous function *c*(*x*−*vt*), which is a solution of Eq. (4). Because *c*(*x*−*vt*) is continuous and Eq. (4) is translationaly invariant, *c*(*x* − *vt* + const) is also a solution that must describe the front propagation from different initial conditions. Given that perturbed and unperturbed systems have slightly different initial conditions their long-time behavior should in general be different by a shift along the *x*-axis. Our simulations, shown in Fig. 8, confirmed that a non-zero phase difference for pushed and pulled waves for small perturbations, but showed a different behavior for locked invasions. Locked fronts returned to their unperturbed positions provided the perturbation did not exceed a certain threshold; otherwise, the position of the front was shifted by an integer number of patches. This difference between locked and unlocked waves is not surprising. The former has a discrete number of distinct steady-state profiles in which the front can settle. Each of these steady-state fronts has a nonzero basin of attraction, so a perturbation must exceed a certain threshold to force the invasion from one steady state into another.

**Figure 8.**
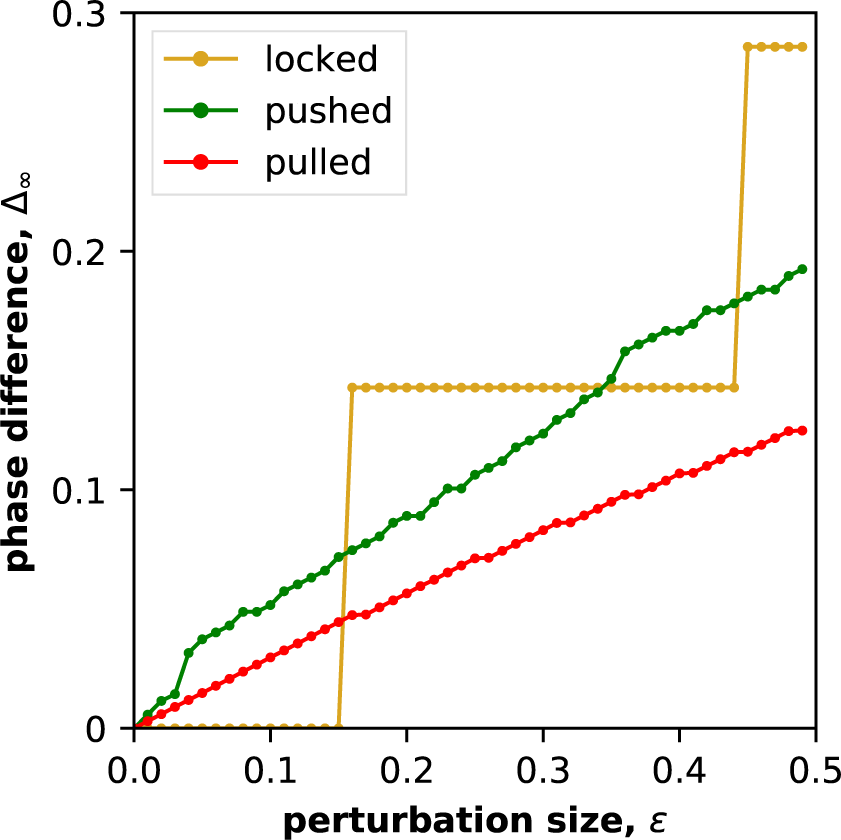
Response to perturbation is different for pulled, pushed and locked waves. The phase difference, Δ_∞_, between the unperturbed and perturbed profiles after 10^5^ time steps as a function of the perturbation strength *ϵ* is shown. We define Δ_∞_ as the number by which the profile of the perturbed wave needs to be shifted back to match the profile of the unperturbed wave. To perform this shift, we determined continuous density profiles as described in Fig. 9. For locked expansions, Δ_∞_ takes discrete values including Δ_∞_ = 0 for weak perturbation. For pushed and pulled waves, Δ_∞_ > 0 provided *ϵ* > 0 and the dependence is continuous. Note that we observe slight undulations in Δ_∞_(*ϵ*) for pushed, but not pulled waves. Parameters for the plots are *c*\* = 0.3 and *K* = 1.0 for the piecewise-linear growth function. The growth and migration rates are *r* = 1.096, *r* = 1.52, *r* = 3.33, and *m* = 0.068, *m* = 0.1432, *m* = 0.20 for locked, pushed, and pulled waves respectively.

There are also differences in how fronts approach the long-time steady state following a perturbation. Exponential relaxation is expected for locked waves, since each of the discrete profiles is a stable fixed point of the dynamics. In contrast, pulled waves are expected to show a much slower relaxation with perturbations decaying as *1/t.* The reasons for this non-exponential relaxation is that the linearized Eq. (4) admits solutions with a continuum of possible velocities and, therefore, its spectrum does not have a gap [39, 59]. Both of these predictions are confirmed by our simulations shown in Fig. 9. We also found that perturbations decay exponentially in time for pushed waves (Fig. 9B). This observations is consistent with the results for CSCT models and is related to the existence of a unique velocity, and, therefore, a spectral gap for pushed waves [59, 71].

**Figure 9.**
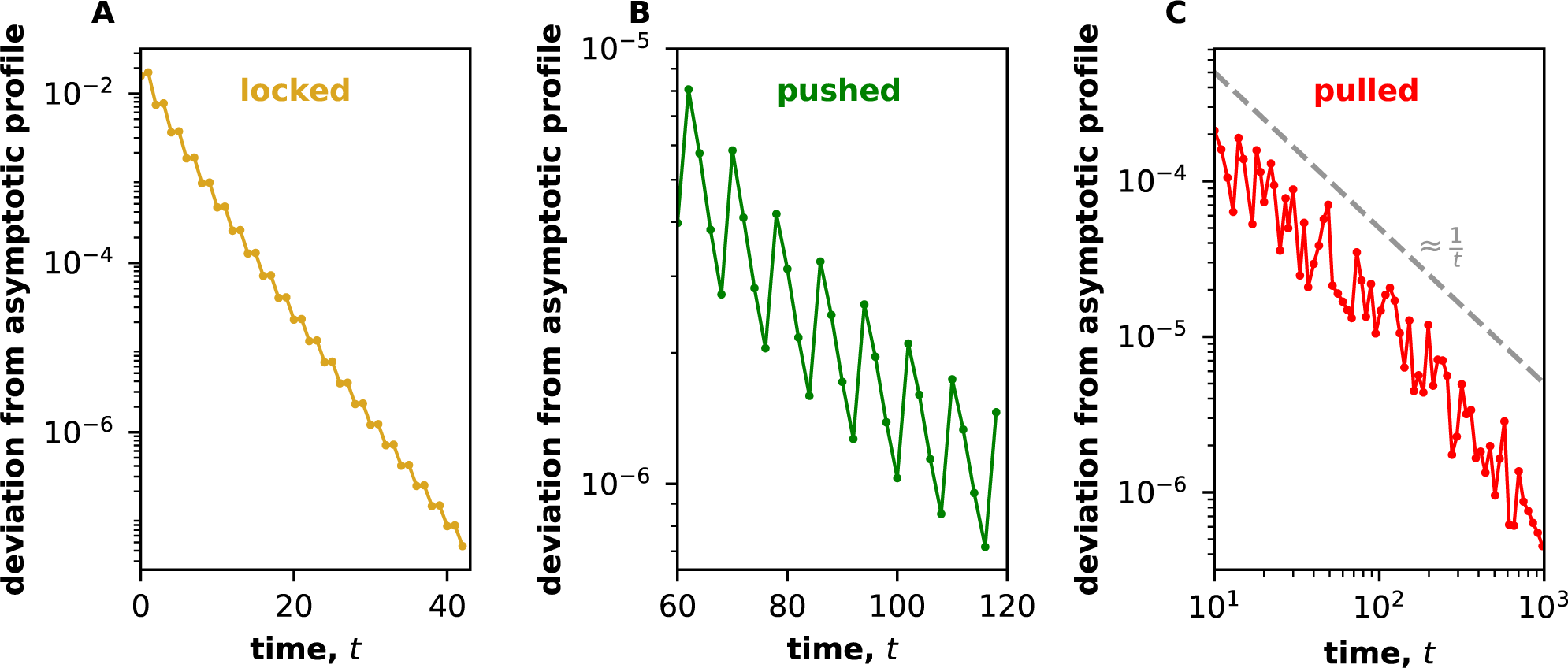
Locked, pushed and pulled invasions show different relaxation dynamics following a perturbation. We show how the front relaxes to its long-time steady state following a perturbation described in text with *ϵ* = 0.75. The asymptotic, steady-state density profile was obtained by first waiting 10^5^ generations for the transient dynamics to die out and then shifting the following 10^3^ profiles in the comoving reference frame. The resulting data was converted into a continuous profile using a cubic spline. The deviation from the asymptotic profile was quantified by the sum of square residuals between the discrete points *c_t_*_,*x*_ and the corresponding points on the continuous steady-state profile. The latter was shifted by a distance that minimized the sum of the square residuals to account for the translational motion of the front. **(A)** and **(B)** Locked and pushed waves showed an exponential decay, which is a straight line on the log-linear plot. **(C)** Deviation for pulled wave decayed as *1/t* which is a straight line on a log-log plot. The thin grey line shows the expected slope of −1. In all panels, *g*(*c*) is the piecewise-linear growth model with *c*\* = 0.3, and *K* = 1.0. The growth and dispersal rates are *r* = 1.1 and *m* = 0.38 for A, *r* = 1.52 and *m* = 0.1432 for B, and *r* = 3.33 and *m* = 0.20 for C.

### Locked invasions are robust to fluctuations

Since locked waves return to exactly the same steady state following a perturbation, they should be robust to demographic and environmental fluctuations. To establish the limits of this robustness, we modified the piecewise-linear growth model and Eq. (4) to include demographic noise, temporal fluctuations of the environment, or spatial heterogeneity of the habitat.

Demographic fluctuations were described by
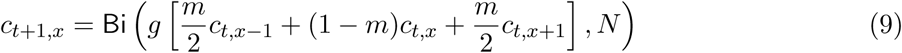

where Bi(*p*, *N*) refers to binomial sampling from *N* trials with the success probability equal to *p.* Temporal fluctuations were modeled by drawing *m* from a uniform distribution from *m* − Δ*m*/2 to *m* + Δ*m*/2 at each time step in Eq. (4). Spatial heterogeneity was included by using a different growth rate in each patch, which was drawn from a uniform distribution between *r* − Δ*r*/2 and *r* + Δ*r*/2.

In all three scenarios, we observed that velocity locking is extremely robust to fluctuations (Fig. 10). Although small plateaus are progressively washed out by stronger fluctuations, large plateaus remain clearly visible even when fluctuations are about 25% of the mean. Thus, velocity locking should be readily observable in experimental and potentially even in natural populations.

**Figure 10.**
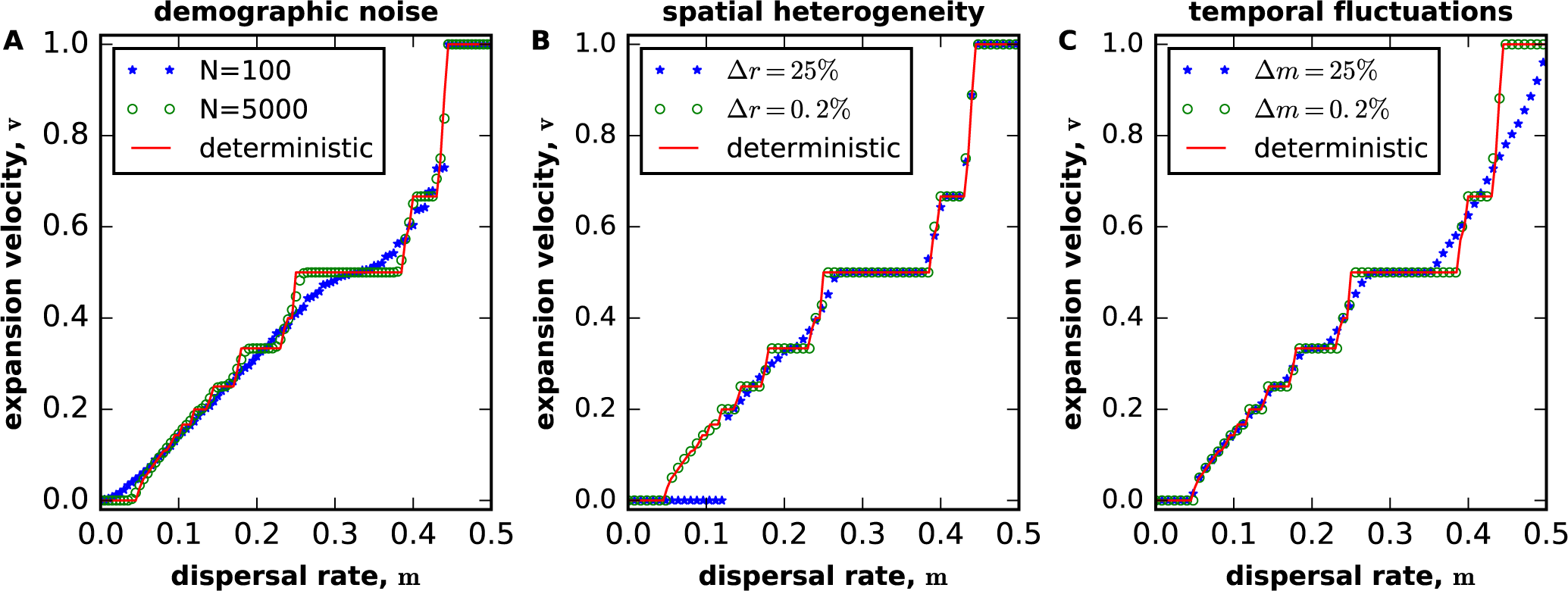
Velocity locking is robust to demographic and environmental fluctuations. Velocity as a function of migration rate is shown in the presence of demographic fluctuations in **(A),** spatial fluctuations in the local growth rate in **(B),** and temporal fluctuations in the global migration rate in **(C).** In all cases, small levels of noise produce clear plateaus as in the deterministic model. Larger levels of noise smooth out small plateaus, but preserve large plateaus. Velocity plateaus disappear completely only for very strong noise. In all panels *r* = 0.93, and *c*\* = 0.22. In B and C, *K* = 1.

### Velocity locking suppresses front diffusion

Given that velocity plateaus remain even in the presence of strong fluctuations, we decided to examine how velocity locking affects stochastic properties of invasion fronts. The consequences of demographic and environmental noise are understood relatively well in CSCT models [39, 72–75]. The main differences between stochastic and deterministic models are fluctuations of the front shape and diffusive wandering of the front position. The latter could be defined in a number of ways, for example, as the integral of population density normalized by the carrying capacity. For an ensemble of independent simulations, the mean position of the front grows linearly in time as in deterministic simulations. The variance of the front position also grows linearly in time as if the front is performing an unbiased random walk relative to its expected position. The effective diffusion constant of the front, *D_f_*, can then be obtained from simulations by fitting the linear increase of the variance to *2D_f_t.* In some cases, the effective diffusion constant of the front can also be computed analytically [39, 72–75].

We applied this procedure of estimating *D _f_* to DSDT models. Figure 11A shows that noisy invasions in DSDT models also exhibit diffusive front wandering and, therefore, can be assigned an effective diffusion constant. We examined how this diffusion constant depends on velocity locking by measuring *D _f_* as a function of *m* for parameter values that result in both locked and unlocked fronts (Fig. 11B). We found that *D _f_* vanishes inside velocities plateaus, but shows very large peaks immediately outside the plateaus. These findings parallel the behavior found in mode locked systems [76], and can be explained by the dynamics without noise. Indeed, Fig. 8 shows that, for locked invasions, small perturbations are quickly forgotten and, ultimately, have no effect on the front position. Therefore, *D _f_* must be zero for locked fronts. For pulled and pushed fronts, however, any perturbations leads to a small shift in the asymptotic position of the front, so repeated perturbations due to demographic or environmental fluctuations must result in a random walk. Hence, *D_f_* > 0 outside velocity plateaus. The peaks in *D _f_* on plateau boundaries are explained by the rapid variation of *v* with *m* in these regions, which results in extreme sensitivity of the front motion to perturbations.

**Figure 11.**
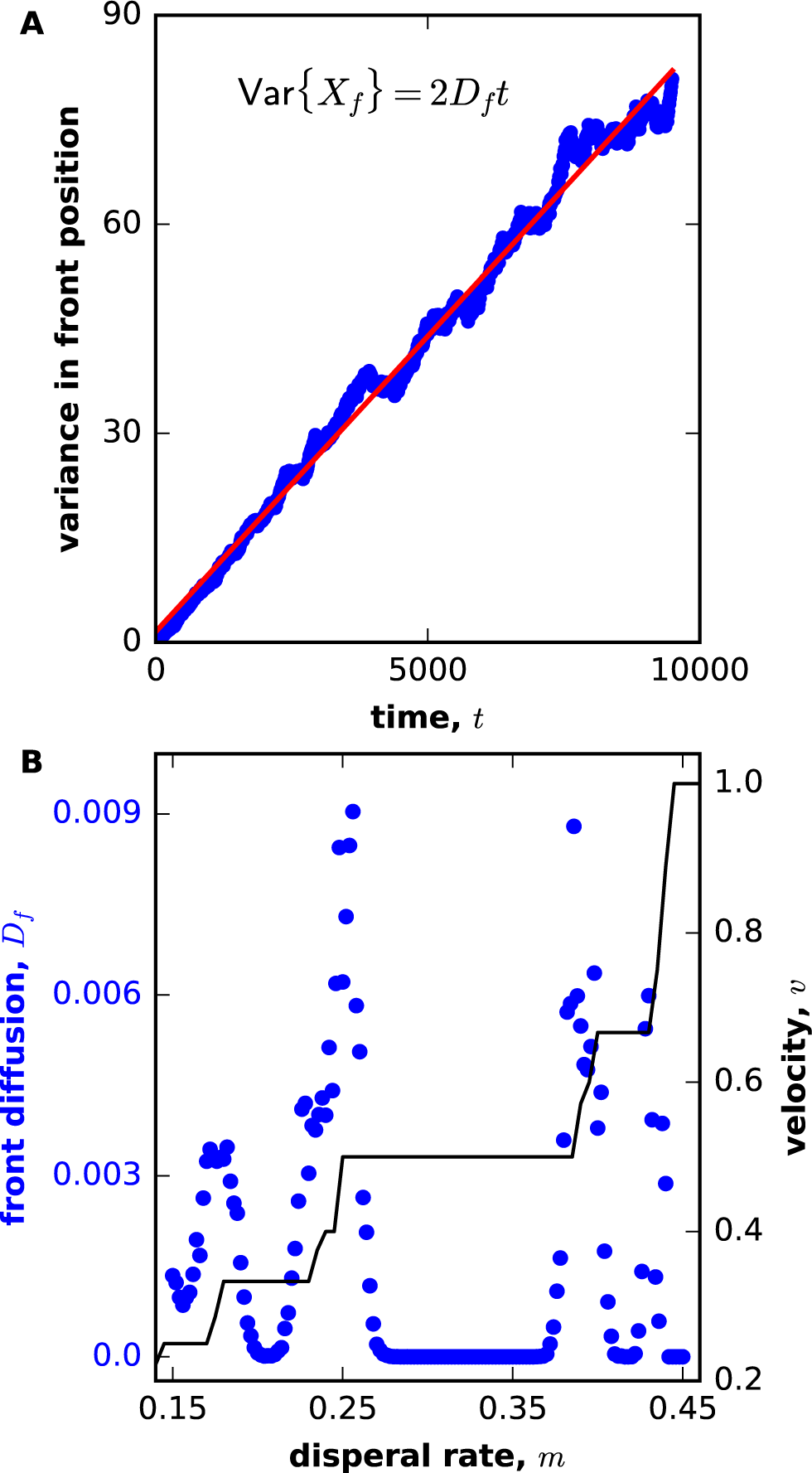
Dramatic changes in front diffusion between locked and unlocked regimes. **(A)** shows that the variance of the front position increases linearly in time and can therefore be used to define effective diffusion constant of the front, *D _f_.* **(B)** The dependence of *D _f_* indicates major differences between locked and unlocked regimes. Inside velocity plateaus *D_f_* = 0, but it is nonzero in unlocked regions. The diffusion constant is especially large near plateau boundaries, where *v*(*m*) changes rapidly. For convenience, *v*(*m*) is plotted in thin black line. Here, we simulated the piecewise-linear model with demographic noise: *r* = 0.93, *c*\* = 0.22, *N* = 2000. To determine each value of *D_f_*, we ran simulations for 9500 generations and averaged over 100 independent runs. The position of the front was measured as the total normalized biomass.

The vanishing of *D _f_* for locked invasions provides a convenient way to detect velocity locking in situations where one cannot modify the model parameters and confirm the existence of a velocity plateau. Zero diffusion could also be beneficial in technological applications that require coherence or reproducibility. In such case, one might prefer to operate the system at one of the velocity plateaus. On the other hand, the anomalously large *D _f_* near plateau boundaries could explain extreme variability of some invasions or could be used to amplify variability in situations where it is beneficial.

### Locked waves due to positive density-dependent dispersal

Mode locking requires nonlinear dynamics. So far, we focused on an Allee effect as the source of this nonlinearity. In the context of range expansions, density-dependent dispersal could provide an alternative mechanism. To test this hypothesis, we modified Eq. (4) as follows
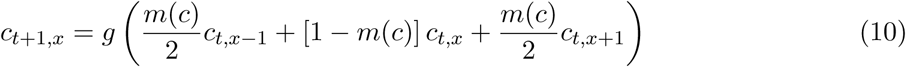

where *m*(*c*) = *m*_0_ + *m*_1_*c* describes the dependence of the dispersal rate on population density. In principle, *g*(*c*) could be arbitrary, but we focus on the piecewise-linear *g*(*c*) with no Allee effect (*rc*\* = *K*) to show that density-dependent dispersal alone is sufficient to produce locked invasions.

For *m*_1_ < 0, dispersal is highest at the edge of the front, and the invasions are pulled. For *m*_1_ *>* 0, the density-dependence is positive, and the linear theory of pulled waves may not apply. Indeed, there are deviations from Eq. (8) and clear velocity plateaus for the sufficiently large ratios of *m*_1_/*m*_0_ (Fig. 12A). The phase diagram in the *m*_0_−*m*_1_ space is qualitatively similar to that in *r*−*m* space, but pushed waves are more prevalent in the *m*_0_ − *m*_1_ space presumably because *m*(*c*) is a continuous function (Fig. 12B).

**Figure 12.**
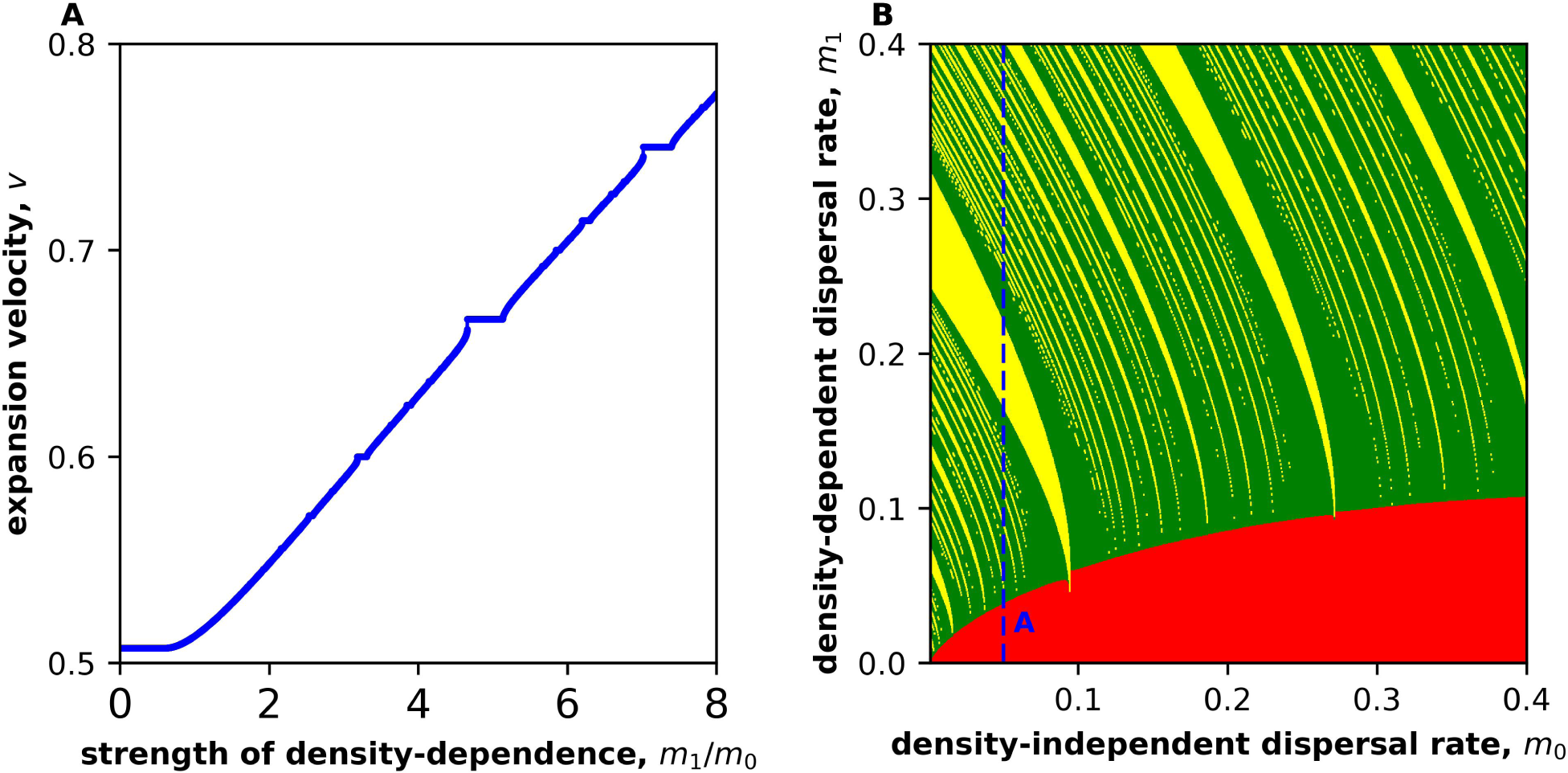
Velocity locking due to positive density-dependent dispersal. **(A)** shows how expansion velocity depends on *m*_1_/*m*_0_, the ratio of density-dependent to density-independent components of dispersal in the model defined in the paragraph below Eq. (10). In simulations, *m*_0_ = 0.05 and *m*_1_ was varied. For small *m*_1_, the velocity is independent of *m*_1_ because the expansions are pulled. For greater *m*_1_, expansions become pushed, which is evident the monotonic and smooth increase of *v* with *m*_1_. At some values of *m*_1_, clear velocity plateaus are observed indicating the existence of locked expansions. **(B)** shows the regions of pulled, pushed, and locked waves in the *m*_0_ − *m*_1_ plane. Note that there are no pinned waves because they require a strong Allee effect. The color scheme and numerical procedures are the same as in Fig. 7A. In both panels, we used a piecewise-linear growth model without an Allee effect: *r* = 3.33, *c*\* = 0.3, and *K* = 1.0.

Density-dependent dispersal has been described in many species, and it can also be easily engineered in microbes [77–79]. Therefore, nonlinearities in dispersal could provide an alternative route to velocity locking in expanding populations.

### Evolution in locked invasions

Given that small changes in model parameters do not affect the velocities of locked fronts, it is not clear whether natural selection can act on mutations that keep the parameters within the Arnold tongue and, therefore, do not change the invasion velocity. We were especially interested in the selection for higher dispersal or growth, which has been repeatedly observed in many species [80–83]. To this purpose, we considered the competition between two genotypes with different dispersal rates (the results for different growth rates were similar).

The competition is described by an extension of Eq. (4) to two genotypes labeled by index *α*; for the resident, *α* = 1 and, for the mutant *α* = 2. This extended equation is most clearly formulated in terms of separate dispersal and growth steps:
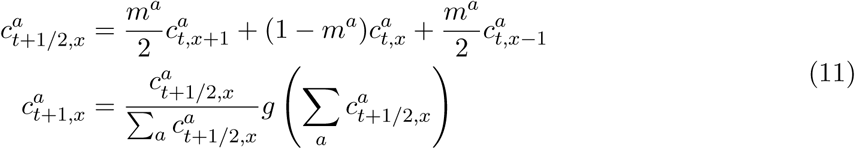

where *t*+1/2 denotes the densities after the dispersal step. Because we assume that the growth rates of the genotypes are the same, the second equation leaves the relative frequencies of the genotypes unchanged. These frequencies do change under the action of the first equation because *m*^1^ ≠ *m*^2^.

We found that dispersal mutants can take over the population even though they have the same invasion velocity as the resident (Fig. 13). Similar to previous findings for CSCT models [40], selection for both faster and slower dispersal was possible depending on the strength of the Allee effect. We therefore conclude that velocity locking does not arrest evolutionary dynamics in the expanding population. Velocity locking can nevertheless lead to unusual dynamics: Dispersal may slowly increase without any appreciable change in velocity until it reaches the plateau boundary, and the velocity suddenly jumps to a different plateau. Such behavior can be easily misinterpreted as an unusual genetic change or strong epistasis.

**Figure 13.**
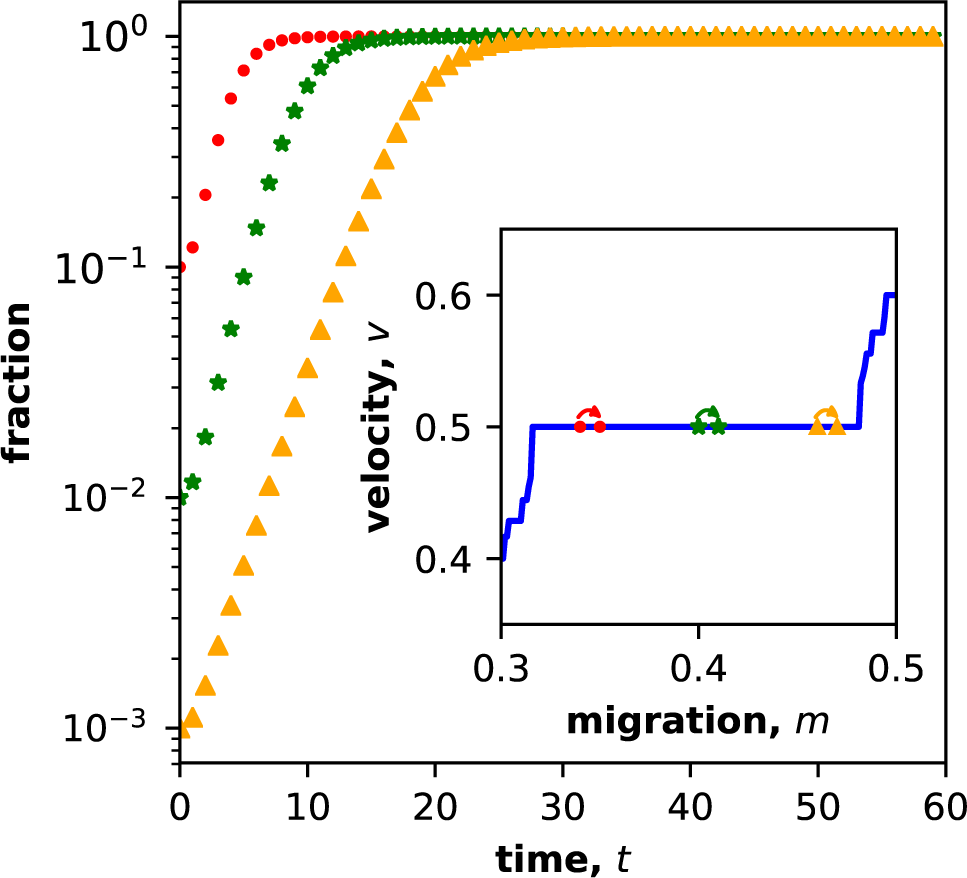
Natural selection occurs despite velocity locking. Mutants with higher dispersal rates invade even though they do not increase the expansion velocity. Three sets of *m* for resident and invaders are chosen from a velocity plateau at *v* = 1/2 (inset). The migration rate of the mutant is 0.01 greater than the resident. We used different initial fractions of the invaders, namely 0.1, 0.01, and 0.001 to improve visual clarity. Here, we used the piecewise-linear growth model with *r* = 1.1, *c*\* = 0.3, and *K* = 1.0.

### Velocity locking in two spatial dimensions

Since many range expansions occur in two rather than in one spatial dimensions, we sought to determine how spatial dimensionality affects velocity locking. To answer this question, we generalized Eq. (4) to two spatial dimensions:
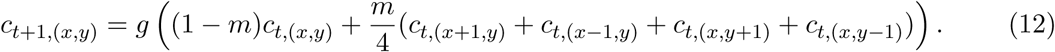

The results of two-dimensional simulations are shown in Fig. 14. We found that velocity plateaus are still present in two spatial dimensions; in addition, the fronts of the expansions assume characteristic geometric shapes that depend on the magnitude of the expansion velocity.

**Figure 14.**
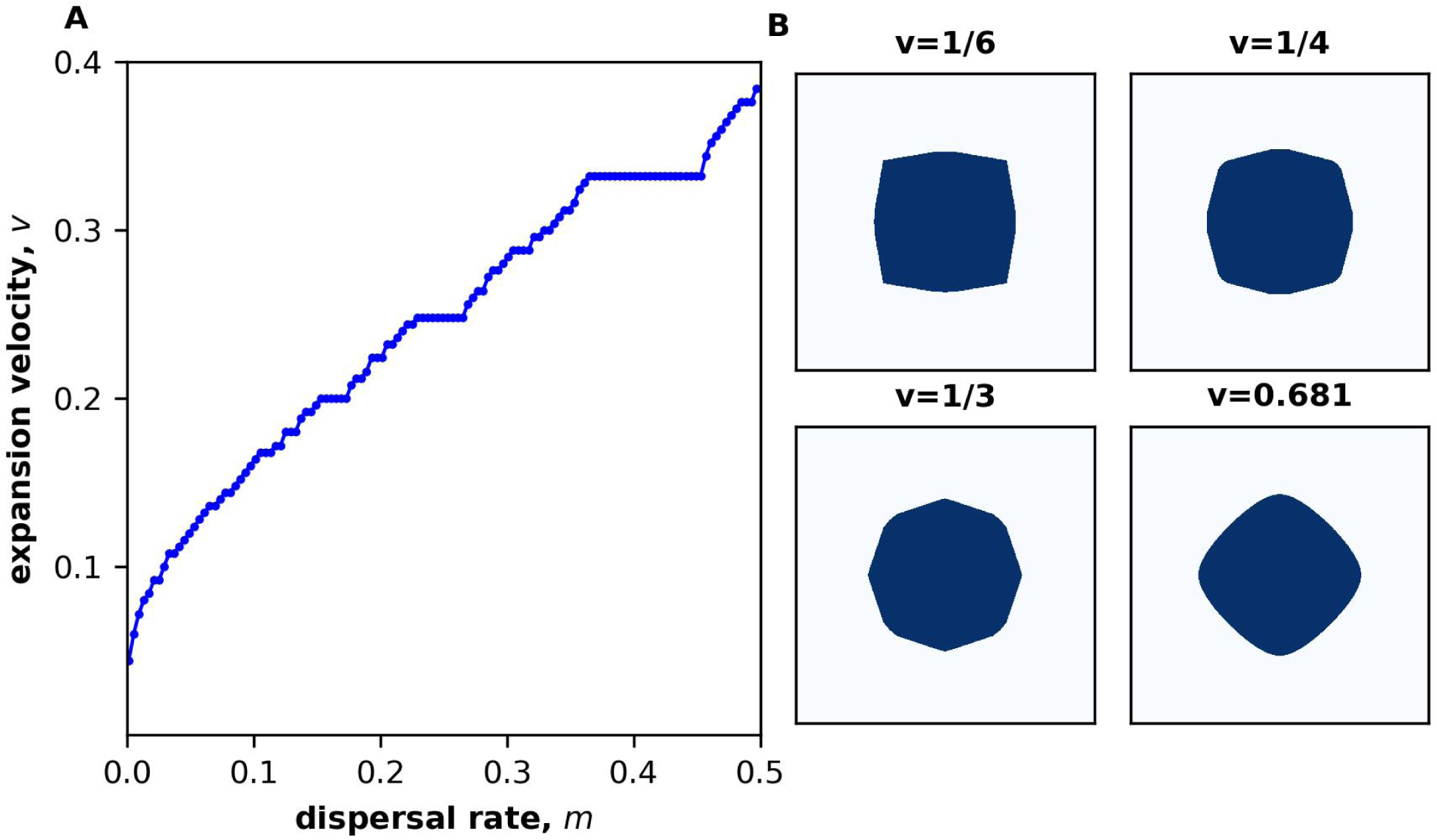
Velocity locking also occurs in two spatial dimensions. **(A)** The dependence of the expansion velocity on the dispersal rate shows clear plateaus. Velocity was measured via a linear regression between the position of the front and the elapsed number of generations. **(B)** shows the invariant shapes of the expanding populations for different expansion velocities. Pulled waves (v = 0.681) have characteristic diamond shape [84–86]. The shapes of locked waves depend on the periodicity of the expansion as shown with three examples. In simulations, *m* = 0.2, and we used the piecewise-linear growth function with *c*\* = 0.3, *K* = 1.0. The growth rates were *r* = 1.2 for A and *r* = 0.95,1.15,1.35 and 3.33 in B. The velocities were calculated by fitting a least-square regression to the front. A static box of 1000 × 1000 patches was used for all simulations in two dimensions.

The results of two-dimensional simulations are shown in Fig. 14. We found that velocity plateaus are still present in two spatial dimensions. In addition, the expansion fronts assume characteristic geometric shapes that depend on the type of the expansion. For pulled waves, the colonized region has the shape of a smoothed rhombus, which was reported previously in the context of growing clusters in the Eden model [84, 85]. For locked waves, we observed distinct shapes (with sharp edges) that depend on the rational numbers *p* and *q* that determine expansion velocity *v* = *p/q* and the periodicity of the population dynamics.

### Velocity locking in CSCT models with spatial and temporal periodicity

Although velocity locking arises most naturally in DSDT models, it is the spatio-temporal periodicity rather than discreteness that is required for velocity locking. To demonstrate this explicitly, we simulated the following CSCT model, where the species experiences seasonal growth and moves with different dispersal rates depending on the type of the traversed patch:
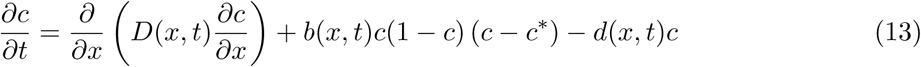

where *D*(*x*, *t*), *b*(*x*, *t*), and *b*(*x*, *t*) are the dispersal, birth, and death rates respectively. We solved Eq. (13) numerically as described in Methods for the following choice of *D*(*x*, *t*), *b*(*x*, *t*), and *b*(*x*, *t*):
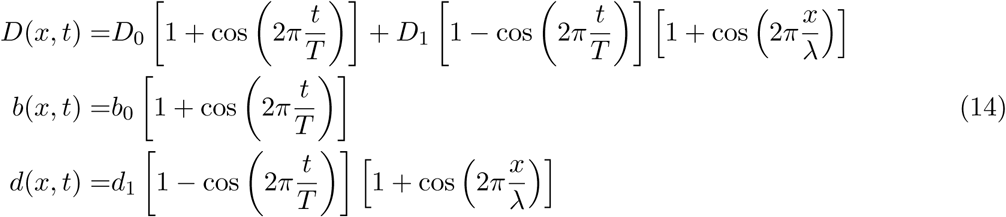

The results are shown in Fig. 15, which confirms the existence of velocity plateaus in CSCT models that possess both spatial and temporal periodicity.

**Figure 15.**
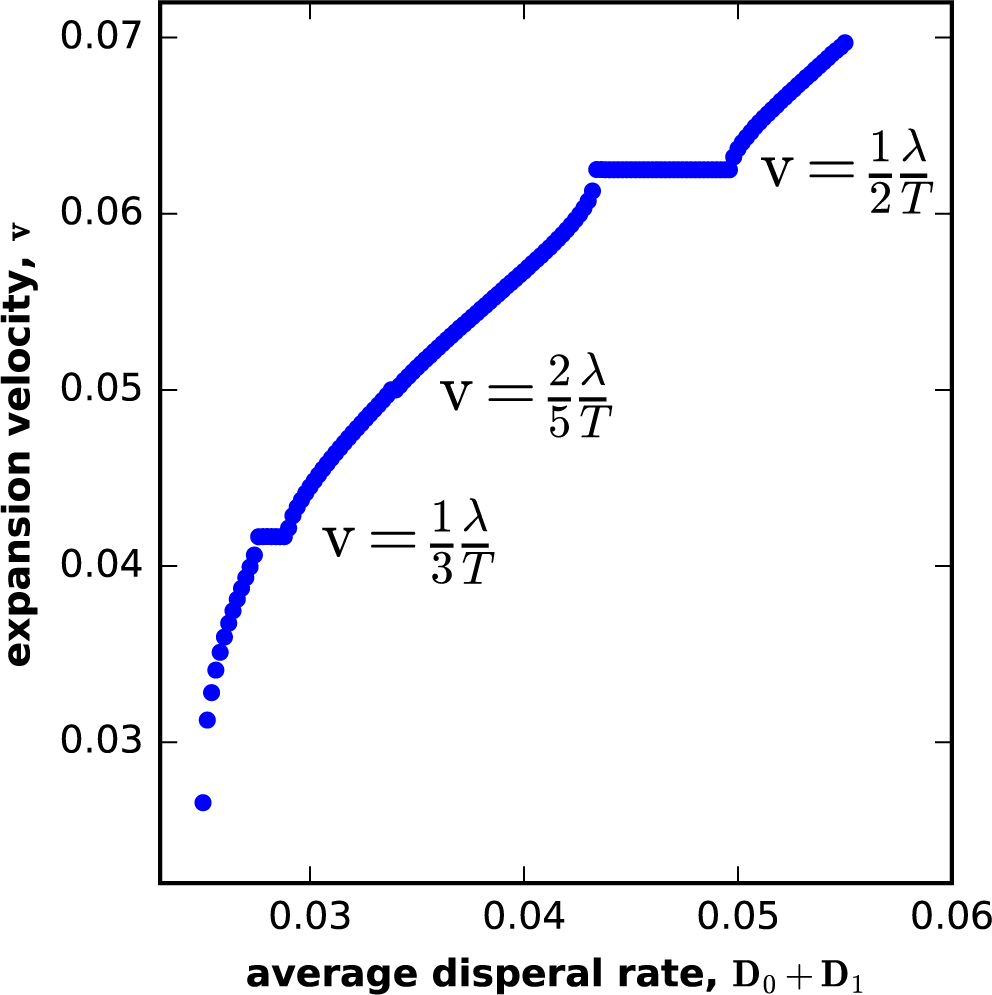
Velocity locking in a CSCT model with spatial and temporal periodicity. The dispersal and growth rates varied in time and space with period *T* and *λ* respectively; see Eqs. (13) and (14). The plot shows both regions of continuous dependence and velocity locking at rational fractions of *λ/T.* Here, *T* = 160, *λ* = 20, *D*_1_ = 0.025, *b*_0_ = 0.25, *d*_1_ = 0.04, *c*\* = −0.2, *D*_0_ is varied. The temporal and spatial average of the dispersal, *D*_0_ + *D*_1_, is plotted on the x-axis. The time step in simulations was Δ*t* = 0.0625 and the spatial time step was set to 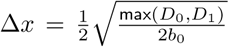 to ensure a stable and accurate discretization. Velocities were computed by a linear fit to the front position in time, where the front position was defined as the location of the furthest site with population greater than half the average carrying capacity.

## Discussion

In this paper, we examined the properties of range expansions in fragmented habitats with seasonal growth. Both ecological conditions are quite common in natural and laboratory populations; for example, in systems with liquid-handling robots, microfluidic devices, regular population arrangements found in agricultural settings, and for species shifting towards more fragmented and seasonal habitats in response to raising temperatures [12, 21–24, 27–31, 31–34, 46, 87]. Invasions under such conditions are better described by models with discrete space and time rather than partial differential equations [36, 37]. Therefore, understanding the effects of discreteness could have important implications for biotechnology and ecosystem management.

Discrete models also arise when continuous models are solved on a computer. Although continuous behavior is typically assured in the limit of infinitely fine discretization,^1^ large discretization steps are often chosen in practice because one is willing to sacrifice accuracy for computational efficiency. The key question is then whether the difference between continuous and discrete models is only quantitative or whether there are qualitative differences between the alternative approaches to model population dynamics.

When both space and time are discrete, range expansions can fundamentally differ from the predictions of continuous models. In particular, invasions can proceed in a step-like or pulsed fashion with population densities at the front assuming a discrete set of values that repeat periodically. Such invasions have been observed in the field although they have been attributed to other factors [88, 89]. The velocities of pulsed expansions are locked and completely insensitive to moderate variations in the rates of dispersal and growth. However, when parameter change exceeds a certain threshold, expansion velocity and the periodicity of front pulsations change discontinuously. These dynamics produce a characteristic pattern of plateaus on the plot of a response variable (velocity) vs. a control variable (dispersal) known as the Devil’s staircase in physics [51]; see Fig. 2.

Why does velocity locking occur only in DSDT models and not for pulled waves? The answer lies in the translational symmetry of the traveling wave solutions [90]. Consider an expansion that has reached a steady state and expands at constant velocity with a time-invariant density profile in the reference frame co-moving with the expansion. When space is continuous, a shift of the density profile by any distance results in a density profile that is also a traveling wave solution of the underlying mathematical model. In consequence, there is no restoring force that would oppose a spatial translation of the invasion front, and any perturbation produces a nonzero shift in the front position. Repeated perturbations then modify the invasion rate. Since changes in the model parameters can be viewed as such perturbations, we expect that models with continuous space cannot exhibit velocity locking.

Invasions in DSCT models also possess translational symmetry because two consecutive density profiles separated by any time are both solutions of the underlying mathematical model. Since time is continuous, there is again a continuous family of solutions, which are also related by spatial translations since density profiles at different times are simply shifted relative to each other. The only exception are pinned invasions because a temporal translation does not result in a distinct density profile when the velocity is zero. As a result, DSCT models exhibit velocity pinning, but not locking for exactly the same reasons as the continuous space models.

For pulled waves, velocity locking does not occur because of another continuous symmetry unrelated to spatial and temporal continuity. The velocity of pulled invasions is determined by the dynamics at the expansion edge, where the population density is small and all nonlinear terms can be neglected. Since linear equations are invariant under multiplication by any positive number, there is a continuous family of solutions, which can be interpreted as a family of temporal or spatial translations. Using the same arguments as for CSCT models, we then conclude that pulled expansions should never experience velocity locking.

This symmetry based analysis predicts that models that do not exhibit velocity locking should have continuous profiles in the comoving reference frame. This prediction agrees with the known results and our simulations. When space is continuous, the density profile is clearly a continuous function, which translates in space as the expansion proceeds [1, 18, 19, 59]. Although the density profile at any given time assumes only a discrete set of values in DSCT models, the density profile is nevertheless continuous in the comoving reference frame when density profiles at different times are shifted relative to each other by the distance that they have traveled [18, 19]. Similarly, pulled and pushed waves in DSDT models have a continuous density profile in the comoving reference frame despite the fact that both space and time are discrete (Fig. 4A and SI). In sharp contrast, the density profile of locked invasions is clearly discrete even in the comoving reference frame (Fig. 5).

The remaining question is why velocity locking occurs in DSDT models with significant nonlinearities in dispersal or growth. An intuitive argument was proposed in Ref. [53] that noticed that the infinite dimensional system of coupled difference equations in DSDT models can be reduced to a single difference equation. This dimensional reduction is based on the time scale separation between the dynamics of front shape and position, which occurs because perturbations decay exponentially fast in invasions that are not pulled (Fig. 9). Thus, the position of the front *X_f_* can be described by the following map
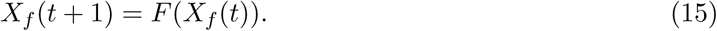

The function *F* is necessarily periodic, so the dynamics can be restricted to *X_f_* ∊ (0, 1) for DSDT models or to *X_f_* ∊ (0, λ) for periodic CSCT models (in this case one considers *X_f_* at times *t* and *t* +*T*). Because of this periodicity *F* is often referred to as a circle map.

Mode locking in circle maps is well understood. For *F* that are monotonic, Eq. (15) predicts an aperiodic increase of *X_f_* similar to the dynamics in pulled waves [51]. However, once nonlinearities are strong enough to make *F* non-monotonic, the circle maps can lock into periodic oscillations. These oscillations arise from the fact that *F_q_*, which is the result of *q* consecutive applications of *F*, develops an attractive fixed point: *F_q_*(*X_f_*) = *X _f_*.^2^ The existence of such fixed points explains the periodicity, stability, and insensitivity to parameter changes of locked waves.

The theory of mode locking in circle maps also predicts that not all parameter values that control the shape of *F* result in a stable fixed point of *F_q_* [51, 51, 67, 68]. In the context of range expansions, this means that velocity plateaus account for only part of the v(m) plot−the Devil’s staircase is not complete. Therefore, at some values of *m*, front propagation occurs without strict periodicity and velocity locking even though the expansions are not pulled. We termed such unlocked expansions pushed waves. Consistent with the above arguments, we found that pushed and locked invasions occur in DSDT models only in the presence of positive density-dependence in dispersal or growth. Negative density-dependence instead produces pulled invasions, which proceed in a smooth, continuous fashion in both discrete and continuous models.

Velocity locking could have important implications for the management of invasive species because of the unusual way in which invaders respond to interventions. Indeed, a management strategy could be deemed ineffective when it does not result in any change of the locked invasion velocity. Yet, a dramatic reduction in the rate of invasion may occur if the management effort is intensified beyond a point necessary to push the population to a new velocity plateau, say from *v* = 1/2 to *v*= 1/4 or even *v*= 0.

Velocity locking also affects the dynamics and evolution of the invading population. We found that locked fronts propagate quasi-deterministically without exhibiting any diffusive wandering due to demographic or environmental fluctuations. In contrast, unlocked fronts have a nonzero effective diffusion constant that becomes especially large near velocity plateaus. While velocity locking does not suppress the evolution of dispersal and growth, it nevertheless leads to important differences compared to the dynamics in continuous models. The analysis of reaction-diffusion equations suggests mutants are often selected on their expansion velocity [40, 91]. For locked invasion, mutations that do not change the expansion velocity can nevertheless be selected (Fig. 13). Sequential accumulation of such mutation can shift the population to a new velocity plateau, which would appear as the case of both very rapid evolution and very strong epistasis.

Our results show that that velocity locking occurs in both one and two spatial dimensions and is extremely robust. It persists despite external perturbations, demographic noise, environmental fluctuations, and habitat heterogeneity. Velocity locking also does not require perfect discreteness of space and time; approximate periodicity in time and space is sufficient. Therefore, we expect that locked expansions should be easily observable in laboratory experiments and possibly in nature. The lack of observations to date might be attributed to the lack of awareness of mode locking phenomena among the scientists who study range expansions. Indeed, mode locking is not widely known outside nonlinear sciences, and the literature on population biology is dominated by continuous models, in which velocity locking cannot occur.

While the main focus of our work has been on biological invasions, velocity locking could also have implications beyond ecology. Indeed, front propagation arises in quantum chromodynamics [92], entanglement spreading [15, 16], chemical kinetics [14, 93], biofilm growth [94], cancer biology [95], and population genetics of spatial [2, 3, 96] and well-mixed [4, 97] populations. In many of these contexts, the spatio-temporal structure of the environment could be discrete or periodic and, therefore, support velocity locking together with all the unusual behaviors associated with locked fronts.

## Methods

### Simulations of CSCT, CSDT, and DSCT models in Fig. 1

For CSCT, we used a cubic polynomial for *g*(*c*), which can describe growth with and without an Allee effect [38, 61, 98]:
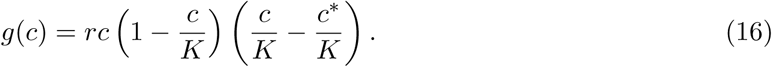

Here, *c*\* controls the strength of an Allee effect, *r* controls the growth rate, and *K* is the carrying capacity. When *c*\* > 0, the Allee effect is strong, and *c*\* is the Allee threshold, i.e. the minimal density required for growth. The Allee effect is weak when −*K* < *c*\* < 0 and absent when *c*\* < −*K.* Simulations were performed using the pdepe function in MatLab^®^, and the discretization was chosen such that the error always met the tolerance of 10^−3^.

For CSDT, we used the a kernel with exponential tails:
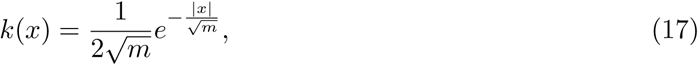

because it leads to a traveling-wave solution with or without an Allee effect. More fat-tailed kernels, e.g. with power-law tails, result in accelerating expansions unless there is a significant Allee effect [18, 19]. For the growth function, we used the piecewise linear model: *g*(*c*) = *rc* for *c* < *c*\* and *g*(*c*) = *K* for *c* ≥ *c^*^.* The simulations performed numerical integration of Eq. (2) using discretization of at least 10^4^ points.

For DSCT models, we used the cubic *g*(*c*) defined in Eq. (16) and solved Eq. (3) for 2000 patches using the standard Runge-Kutta method in MatLab^®^.

In all models, the habitat was empty initially, but the boundary conditions were chosen to mimic an expansion from a region where the population is well established. Specifically, population density was set to the carrying capacity on the left edge of the habitat. The boundary condition on the right edge was absorbing, but we did not run simulations until that boundary was reached. In all simulations, the velocity was obtained via least-square fitting of the front position to a linear function of time.

In Figure 1 we used the following parameters. In A, we used the following parameters. For CSCT, *m* = 3, *r* = 1, *c*\* = 0.25, and *K* = 1. For DSCT, *m* = 2, *r* = 1, *c*\* = 0.25, and *K* = 1. For CSDT, *m* = 0.0204, *r* = 0.2, *c*\* = 0.3, and *K* = 1. For DSDT case the growth was piecewise linear with *m* = 0.25, *r* = 0.7, *c*\* = 0.15, *K* = 1. In B, we used the following parameters. For CSCT, *r* = 4, *c*\* = 0.4, and *K* = 1 for the strong Allee effect and *r* = 0.25, *c*\* = −1.2, and *K* = 1 for no Allee effect. For DSCT, *r* = 1.1, *c*\* = 0.25, and *K* = 1 for the strong Allee effect and *r* = 1.1, *c*\* = −1.1, and *K* = 1 for no Allee effect. For DSCT, *r* = 0.5, *c*\* = 0.3, and *K* = 1 for the strong Allee effect and *r* = 3.3, *c*\* = 0.3, and *K* = 1 for no Allee effect. For DSDT, Beverton-Holt model was used with *A* = 4.1, *B* = 0.3, and *c*\* = 0.2 for the strong Allee effect and with *A* = 4.1, *B* = 1, and *c*\* = 0 for no Allee effect.

### Simulations of DSDT models

Simulations were performed as follows. Each generation, population densities were updated according to Eq. (4) with reflecting boundary conditions on both ends of the simulation box. The size of the simulation box contained at least 100 patches, and its position was periodically adjusted to ensure that the expansion front is centered. Prior to measuring the invasion velocities, we allowed the population to expand for at least 10^4^ generations starting from a step-like initial condition. The velocities were then determined from a least-square linear regression of the position of the population front on time, during a period of at least another 10^4^ generations.

To avoid numerical artifacts associated with uncontrolled machine precision in python, we imposed a cutoff in the density for stochastic simulations at *c* = 10^−^8; thus the any values of *c* < 10^−^8 were set to 0.

### Simulations of CSCT models with spatial and temporal periodicity in Fig. 15

To handle the time and space dependent coefficients in Eq. (13), simulations were performed with the following discretization scheme
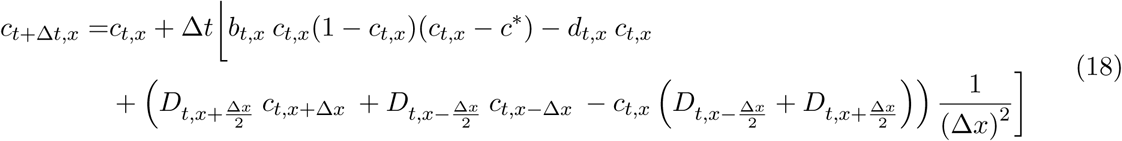

where Δ*t* and Δ*x* were the time and space discretizations used for the numerical solution. By *x* f Δ*x*/2, we indicate the fact that the diffusion constant was evaluated at a spatial position half-way between the discretization points using the analytical form of *D*(*t, x*) in the continuous formulation of the partial differential equation.

Simulations were performed at two separate discretization values to ensure that results were independent of the choice of discretization. To avoid numerical artifacts associated with uncontrolled machine precision, we imposed a cutoff in the density at *c* = 10^−^8; thus the any values of *c* < 10^−^8 were set to 0.

### Determining velocities in simulations

For the simulations of DSCT, CSDT, DSDT and CSCT models; and stochastic simulations; in Figs. 1, 2, 4, 5, 6, 10 and 11 the velocity was measured by a least-square linear regression on the total biomass normalized by the carrying capacity as a function of time. The position of the front in Fig. 11 was measured as the total normalized biomass. Measuring it in this fashion allowed accurate estimation of the front position as a real number.

For the CSCT model with spatial and temporal periodicity in Fig. 15, the velocity was determined by a least-square linear regression of the position of the population front in time at 10 equally spaced points. To define the position of the front, we defined the average carrying capacity of the population, *K*_avg_, as the mean population size far behind the expansion front at that time. The position of the front was then defined as the the furthest position on the x-axis at which the population density was greater than *K*/2.

For the large runs in Figs. 7 and 12 the velocity was measured using 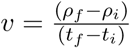, where *ρ _f_* and *t _f_* are total biomass and time at the end and *ρ_i_* and *t_i_* are the initial biomass and time, after a transient of 10^6^ steps or longer. For long transients the velocity is consistent with the least-square linear regression described previously.

In Fig. 14, the velocity in the two dimensional system, was measured by fitting a regression to the front along the x-axis. The front corresponded to the first site where the density was below 0.95K, where *K* is the carrying capacity.

### Velocities of pulled expansions

Here, we derive the velocity of pulled expansions given the low-density growth rate *r* and migration rate *m* in DSDT models. Our calculation closely follows that of van Saarloos [59] and is given here for completeness.

When expansions are pulled, their properties are determined by the dynamics at the expansion front, where *c_t_*_,*x*_ is small and *g*(*c*) can be approximated as *rc.* In the absence of an Allee effect, this approximation yields the maximal possible growth rate; therefore, the bulk dynamics cannot push the population to expand at a higher rate than predicted by the linear approximation. For a weak Allee effect, the magnitude of the growth rate increase in the bulk determines whether the expansion is driven by its edge (pulled waves) or its bulk (pushed waves). For a strong Allee effect, the linear expansion results in *r* < 0, and the linear analysis is clearly unable to describe the expansion dynamics.

The linearization of Eq. (4) yields
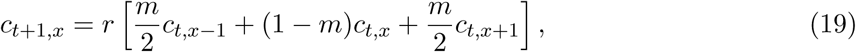

This equation is solved through Fourier transforms defined as
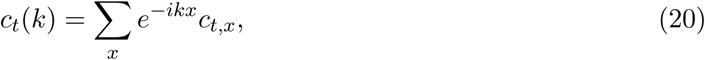

with the following result
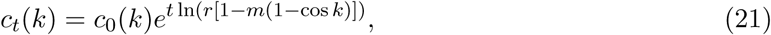

where *c*_0_(*k*) is the Fourier transform of the initial population density.

We then shift to the comoving reference frame with a spatial coordinate *z* = *x* − *vt* and perform the inverse Fourier transform:
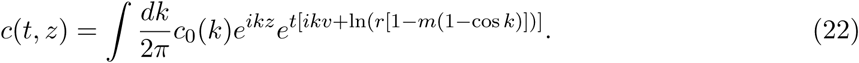

This integral can be evaluated using the saddle point approximation [99], which is appropriate for large *t* when the transient dynamics are over. The saddle point approximation states that the integral is dominated by a region near *k*\*, a particular value of *k* in the complex plane for which the complex derivative with respect to *k* of the power in the second exponent in Eq. (22) is zero:
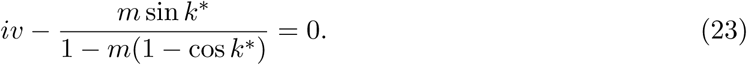

Since this is a complex equation, both real and imaginary parts need to be zero, which yields two real equations for the three unknowns: the real and imaginary parts of *k*\* and *v.* The third needed equations comes from the requirement that *c*(*t, z*) does not increase to infinity or diminish to zero at long times, which is due to the definition of a comoving reference frame. We then obtain the following condition:
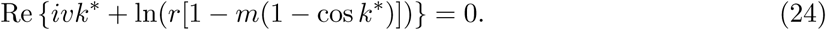

It is easy to see that Eqs. (23) and (24) are satisfied only by purely imaginary *k*\* = *iκ*, which yields the expected exponential decay of *c*(*z*) with *z* as *e*^−*κz*^ from the first exponent in Eq. (22). The velocity *v* and asymptotic decay rate *r.* are then given by the following system of equations.
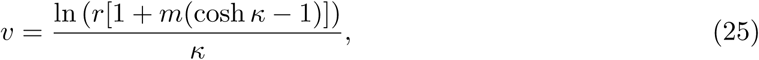

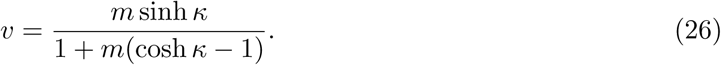

The second equation is equivalent to the requirement that *κ* minimizes *v* in the first equation. Thus, we can alternatively express *v* as
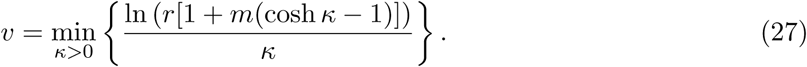

### Expansions locked at 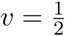 in the piecewise linear model

We now show that *v*(*m*) indeed exhibits exact plateaus by explicitly finding traveling-wave solutions of Eq. (4) with *v* = 1/2 for the piecewise linear model. Since velocity locking occurs only in the presence of an Allee effect, we will assume that *rc*\* < *K.* Our strategy is analogous to that deployed for partial differential equations with piecewise linear growth [41]: We solve in each region of *c* where *g*(*c*) is linear and then match the solutions. For *c* > *c*\*, the solution is *c_t,x_* = *K*, i.e. the bulk of the wave is at the carrying capacity. For *c* < *c*\*, the dynamics are described by Eq. (19), which admits exponential solutions of the form
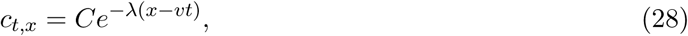

where *C* is the amplitude of the solution that should be determined from matching to the solution behind the front, and *λ* is the spatial decay rate chosen to satisfy Eq. (19) with *v* = 1/2, i.e. we require that
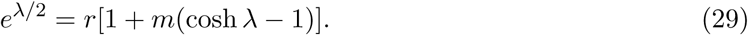

To analyze the nature of solutions to this equation, it is convenient to define *y* = *e^λ^*/^2^ − 1 and rewrite the equation as follows:
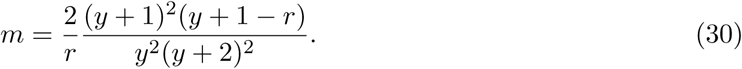

For *r* ≤ 1, i.e. for the strong Allee effect, the right hand side is monotonically decaying from +∞ to 0 as *y* increases from its minimal value of 0 to +∞. As a result, there is a unique solution *λ* (*m*) of Eq. (29). When the Allee effect is weak (*r* > 1), the right hand side of equation Eq. (30) increases from − ∞. to a maximum (*m_e_*) and then declines to 0 as *y* increases from 0 to +∞. Hence, there may be two solutions when the migration rate is below that maximum (*m* < *m_e_*) or no solutions when it is above the maximum (*m* > *m_e_*). The value of *m_e_* is defined as
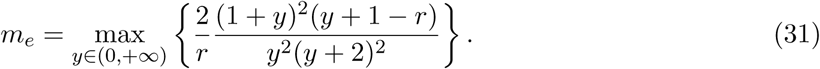

Clearly, for *r* > 1, Eq. (31) sets an upper bound on the migration rates consistent with *v* = 1/2. We also note that, when Eq. (29) has two solutions, only the larger one corresponds to a pushed expansion. This situation is completely analogous to that for reaction-diffusion equations discussed in Ref. [59]. The smaller *λ* corresponds to solutions arising from initial conditions that decay very slowly at +∞ and therefore lack biological realism since all expansions are started by a population completely confined to a finite region of space.

With *λ* defined by the largest solution of Eq. (29), the solution for *c_t,x_* is then given by the following ansatz:
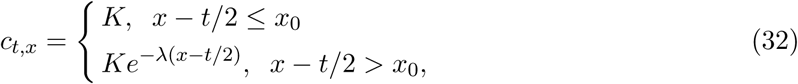

where *x*_0_ determines the initial position of the front. In the following, we assume that *x*_0_ = 0 for simplicity. Note that we set *C* = *K* due to the matching requirement. Indeed, for the exponential profile to be preserved in the nonlinear model, the right most point of the bulk solution must also satisfy Eq. (29) in the generation following the update that transforms the left most point of the front solution to the right most point of the bulk solution. In other words, *C* = *K* ensures that the density of the left most point below the carrying capacity is always consistent with the exponential decay at the front.

Three other conditions are necessary for the ansatz to hold. (i) The right most point of the bulk region should not fall below *c*\* following migration; otherwise, it will fall below the carrying capacity and would not be able to send enough migrants to its neighbor that needs to reach the carrying capacity in the next generation. (ii) The density of the first point below the carrying capacity must not increase above *c*\* after migration following the generations when *x* − *t/2* is an integer; otherwise the bulk region will advance by one step every generation instead of every other generation. (iii) The opposite must be true following the generations when *x* − *t*/2 is a half integer to ensure that the bulk region advances by one patch every two generations. Below we state these conditions as appropriate inequalities and identify the region of *m* ∊ (*m*_min_, *m*_max_) that is consistent with *v* = 1/2, i.e. we compute the location of the corresponding velocity plateau.

First condition:
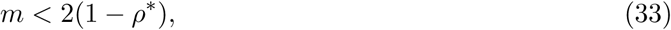

where we defined *ρ*\* = *c*\*/*K*.

Second condition:
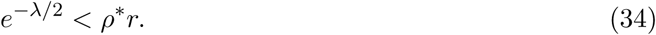

Third condition:
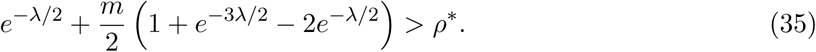

The last two conditions can be used to obtain analytical expressions for *m*_min_ and *m*_max_ by changing inequalities to equalities, which are satisfied at these “critical” values of *m*, using Eq. (29), performing a change of variables *l* = *e*^−λ/2^, and solving for the critical values of *l.* Some of these expression, however, are sufficiently complex and hard to gain intuition from; moreover, one still needs to ensure that the critical values of *l* correspond to the largest solution of Eq. (29). In consequence, it is often much more straightforward to find λ(*m*) numerically from Eq. (29) and verify that all three conditions are satisfied.

For completeness, we also provide the solutions for the critical values of *m* discussed above:
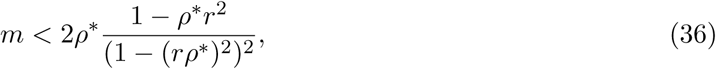

which is applicable when λ = −2ln(*ρ*\**r*) is the largest root of Eq. (29),
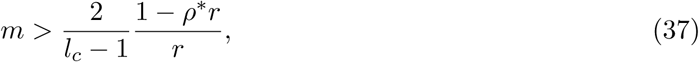

which is applicable when λ = 2ln(*l_c_*) is the largest root of Eq. (29), and *l*, is the positive root of
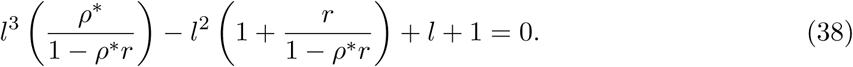

When λ = 2ln(*l_c_*) is the only or the smallest root of Eq. (29), there is no velocity locking.

Note that in addition to Eq. (36), *m* is bounded above by *m_e_* when *r* > 1 and the requirement that *m* ≤ 0.5 in our simulations.

## Acknowledgements

Preliminary work was carried out by Vipul Vachharajani, Eugene Yurtsev, and Jeff Gore, who observed velocity locking in simulations. This work was partially supported by a grant from the Simons Foundation (#409704, Kirill S. Korolev), by the Cottrell Scholar Award (#24010, Kirill S. Korolev), and by a grant from Moore foundation (#6790.08, Kirill S. Korolev). Simulations were carried out on Shared Computing Cluster at Boston University.

## Footnotes

1 The transition from DSDT to CSCT models and the associated smoothing of velocity plateaus were studied in Ref. [53]

2 non-monotonicity of *F* is required for an attracting fixed point in the *F_q_* because any iterate of a monotonic *F* is monotonic.

## References

[1] Murray, J. D. Mathematical Biology (Springer, 2003).

[2] Fisher, R. A. The wave of advance of advantageous genes. Annals of Eugenics 7, 355–369 (1937).

[3] Kolmogorov, A. N., Petrovsky, N. & Piscounov, N. S. A study of the equation of diffusion with increase in the quantity of matter, and its application to a biological problem. Moscow University Bulletin of Mathematics 1, 1 (1937).

[4] Tsimring, L. S., Levine, H. & Kessler, D. A. Rna virus evolution via a fitness-space model. Physical review letters 76, 4440 (1996).

[5] Brockmann, D. & Helbing, D. The hidden geometry of complex, network-driven contagion phenomena. Science 342, 1337–1342 (2013).

[6] Pateman, R. M., Hill, J. K., Roy, D. B., Fox, R. & Thomas, C. D. Temperature-dependent alterations in host use drive rapid range expansion in a butterfly. Science 336, 1028–1030 (2012).

[7] Phillips, B. L., Brown, G. P., Greenlees, M., Webb, J. K. & Shine, R. Rapid expansion of the cane toad (bufo marinus) invasion front in tropical australia. Austral Ecology 32, 169–176 (2007).

[8] Ramaswamy, S., Toner, J. & Prost, J. Nonequilibrium fluctuations, traveling waves, and instabilities in active membranes. Physical review letters 84, 3494 (2000).

[9] Weiner, O. D., Marganski, W. A., Wu, L. F., Altschuler, S. J. & Kirschner, M. W. An actin-based wave generator organizes cell motility. PLoS Biol 5, e221 (2007).

[10] Takamatsu, T. & Wier, W. Calcium waves in mammalian heart: quantification of origin, magnitude, waveform, and velocity. The FASEB Journal 4, 1519–1525 (1990).

[11] Nelson, P. Biological physics (WH Freeman New York, 2004).

[12] Bär, M., Falcke, M., Levine, H. & Tsimring, L. S. Discrete stochastic modeling of calcium channel dynamics. Physical Review Letters 84, 5664 (2000).

[13] Ishihara, K., Korolev, K. S. & Mitchison, T. J. Physical basis of large microtubule aster growth. Elife 5 (2016).

[14] Pelcé, P. & Libchaber, A. Dynamics of curved fronts (Elsevier, 2012).

[15] Schachenmayer, J., Lanyon, B., Roos, C. & Daley, A. Entanglement growth in quench dynamics with variable range interactions. Physical Review X 3, 031015 (2013).

[16] Jurcevic, P. et al. Quasiparticle engineering and entanglement propagation in a quantum many-body system. Nature 511, 202 (2014).

[17] F’ath, G. Propagation failure of traveling waves in a discrete bistable medium. Physica D: Nonlinear Phenomena 116, 176–190 (1998).

[18] Kot, M., Lewis, M. A. & van den Driessche, P. Dispersal data and the spread of invading organisms. Ecology 77, 2027–2042 (1996).

[19] Wang, M.-H., Kot, M. & Neubert, M. G. Integrodifference equations, allee effects, and invasions. Journal of Mathematical Biology 44, 150–168 (2002).

[20] Hallatschek, O. & Fisher, D. S. Acceleration of evolutionary spread by long-range dispersal. Proceedings of the National Academy of Sciences 111, E4911–E4919 (2014).

[21] Abraham, E. R. The generation of plankton patchiness by turbulent stirring. Nature 391, 577 (1998).

[22] Pigolotti, S., Benzi, R., Jensen, M. H. & Nelson, D. R. Population genetics in compressible flows. Physical review letters 108, 128102 (2012).

[23] Wilson, W., Morris, W. & Bronstein, J. Coexistence of mutualists and exploiters on spatial landscapes. Ecological monographs 73, 397–413 (2003).

[24] Kéfi, S. et al. Spatial vegetation patterns and imminent desertification in mediterranean arid ecosystems. Nature 449, 213–217 (2007).

[25] Korolev, K. S. & Nelson, D. Competition and Cooperation in One-Dimensional Stepping-Stone Models. Physical Review Letters 107, 088103 (2011).

[26] Menon, R. & Korolev, K. S. Public good diffusion limits microbial mutualism. Physical review letters 114, 168102 (2015).

[27] Fahrig, L. Effects of habitat fragmentation on biodiversity. Annual review of ecology, evolution, and systematics 487–515 (2003).

[28] Datta, M. S., Korolev, K. S., Cvijovic, I., Dudley, C. & Gore, J. Range expansion promotes cooperation in an experimental microbial metapopulation. Proceedings of the National Academy of Sciences 110, 7354–7359 (2013).

[29] Gandhi, S. R., Yurtsev, E. A., Korolev, K. S. & Gore, J. Range expansions transition from pulled to pushed waves as growth becomes more cooperative in an experimental microbial population. Proceedings of the National Academy of Sciences 113, 69226927 (2016).

[30] Dai, L., Korolev, K. S. & Gore, J. Slower recovery in space before collapse of connected populations. Nature 496, 355–358 (2013).

[31] Zhang, Q. et al. Acceleration of emergence of bacterial antibiotic resistance in connected microenvironments. Science 333, 1764–1767 (2011).

[32] Bechinger, C. et al. Active particles in complex and crowded environments. Reviews of Modern Physics 88, 045006 (2016).

[33] Wioland, H., Woodhouse, F. G., Dunkel, J. & Goldstein, R. E. Ferromagnetic and antiferromagnetic order in bacterial vortex lattices. Nature physics 12, 341 (2016).

[34] Paoletti, M. & Solomon, T. Experimental studies of front propagation and mode-locking in an advection-reaction-diffusion system. EPL (Europhysics Letters) 69, 819 (2005).

[35] Petrovskii, S. V. & Li, B.-L. Exactly solvable models of biological invasion (CRC Press, 2005).

[36] Mistro, D. C., Rodrigues, L. A. D. & Petrovskii, S. Spatiotemporal complexity of biological invasion in a space-and time-discrete predator–prey system with the strong allee effect. Ecological Complexity 9, 16–32 (2012).

[37] de Camino-Beck, T. & Lewis, M. Invasion with stage-structured coupled map lattices: Application to the spread of scentless chamomile. Ecological Modelling 220, 3394–3403 (2009).

[38] Fife, P. C. & McLeod, J. B. The approach of solutions of nonlinear diffusion equations to travelling front solutions. Archive for Rational Mechanics and Analysis 65, 335–361 (1977).

[39] Birzu, G., Hallatschek, O. & Korolev, K. S. Fluctuations uncover a distinct class of traveling waves. Proceedings of the National Academy of Sciences 201715737 (2018).

[40] Korolev, K. S. Evolution arrests invasions of cooperative populations. Physical Review Letters 115, 208104 (2015).

[41] Korolev, K. S. The fate of cooperation during range expansions. PLoS computational biology 9, e1002994 (2013).

[42] Hastings, A. et al. The spatial spread of invasions: new developments in theory and evidence. Ecology Letters 8, 91–101 (2005).

[43] Roques, L., Garnier, J., Hamel, F. & Klein, E. K. Allee effect promotes diversity in traveling waves of colonization. Proceedings of the National Academy of Sciences 109, 8828–8833 (2012).

[44] Kimura, M. & Weiss, G. H. The Stepping Stone Model of Population Structure and the Decrease of Genetic Correlation with Distance. Genetics 49, 561–576 (1964).

[45] Korolev, K. S., Avlund, M., Hallatschek, O. & Nelson, D. R. Genetic demixing and evolution in linear stepping stone models. Reviews of modern physics 82, 1691–1718 (2010).

[46] Keitt, T. H., Lewis, M. A. & Holt, R. D. Allee effects, invasion pinning, and species borders. The American Naturalist 157, 203–216 (2001).

[47] Basler, M., Krech, W. & Platov, K. Y. Theory of phase locking in small j osephson-j unction cells. Physical Review B 52, 7504 (1995).

[48] Maselko, J. & Swinney, H. L. A complex transition sequence in the belousov-zhabotinskii reaction. Physica Scripta 1985, 35 (1985).

[49] Lan, Z., Minář, J., Levi, E., Li, W. & Lesanovsky, I. Emergent devils staircase without particle-hole symmetry in rydberg quantum gases with competing attractive and repulsive interactions. Physical review letters 115, 203001 (2015).

[50] Komura, Y. & Kitano, Y. Long-period stacking variants and their electron-concentration dependence in the mg-base friauf–laves phases. Acta Crystallographica Section B: Structural Crystallography and Crystal Chemistry 33, 2496–2501 (1977).

[51] Bak, P. The devil’s staircase. Physics Today 39, 38–45 (1986).

[52] Carretero-González, R., Arrowsmith, D. & Vivaldi, F. Mode-locking in coupled map lattices. Physica D: Nonlinear Phenomena 103, 381–403 (1997).

[53] Carretero-Gonzalez, R., Arrowsmith, D. & Vivaldi, F. One-dimensional dynamics for traveling fronts in coupled map lattices. Physical Review E 61, 1329 (2000).

[54] Fernandez, B. & Raymond, L. Propagating fronts in a bistable coupled map lattice. Journal of statistical physics 86, 337–350 (1997).

[55] Coutinho, R. & Fernandez, B. Fronts and interfaces in bistable extended mappings. Nonlinearity 11, 1407 (1998).

[56] Coutinho, R. & Fernandez, B. Fronts in extended systems of bistable maps coupled via convolutions. Nonlinearity 17, 23 (2003).

[57] Coutinho, R. & Fernandez, B. Spatially extended monotone mappings. In Dynamics of Coupled Map Lattices and of Related Spatially Extended Systems, 265–284 (Springer, 2005).

[58] TurzIK, D. & Dubcová, M. Stability of steady state and traveling waves solutions in coupled map lattices. International Journal of Bifurcation and Chaos 18, 219–225 (2008).

[59] Van Saarloos, W. Front propagation into unstable states. Physics Reports 386, 29–222 (2003).

[60] Allee, W. & Bowen, E. Studies in animal aggregations: mass protection against colloidal silver among goldfishes. Journal of Experimental Zoology 61, 185–207 (1932).

[61] Courchamp, F., Clutton-Brock, T. & Grenfell, B. Inverse density dependence and the Allee effect. Trends in Ecology & Evolution 14, 405–410 (1999).

[62] Beverton, R. J. & Holt, S. J. On the dynamics of exploited fish populations, vol. 11 (Springer Science & Business Media, 2012).

[63] Chen, D., Irvine, J. & Cass, A. Incorporating allee effects in fish stock recruitment models and applications for determining reference points. Canadian Journal of Fisheries and Aquatic Sciences 59, 242–249 (2002).

[64] Hill, A. V. The possible effects of the aggregation of the molecules of haemoglobin on its dissociation curves. J Physiol (Lond) 40, 4–7 (1910).

[65] Yakubu, A.-A. Allee effects in a discrete-time sis epidemic model with infected newborns. Journal of Difference Equations and Applications 13, 341–356 (2007).

[66] Lui, R. Biological growth and spread modeled by systems of recursions. i. mathematical theory. Mathematical Biosciences 93, 269–295 (1989).

[67] Ecke, R. E., Farmer, J. D. & Umberger, D. K. Scaling of the arnold tongues. Nonlinearity 2, 175 (1989). URL http://stacks.iop.org/0951-7715/2/i=2/a=001.

[68] Umberger, D. K. & Farmer, J. D. Fat fractals on the energy surface. Phys. Rev. Lett. 55, 661–664 (1985). URL https://link.aps.org/doi/10.1103/PhysRevLett.55.661.

[69] Eykholt, R. & Umberger, D. Relating the various scaling exponents used to characterize fat fractals in nonlinear dynamical systems. Physica D: Nonlinear Phenomena 30, 43–60 (1988).

[70] Arnol’d, V. I. Remarks on the perturbation theory for problems of mathieu type. Russian Mathematical Surveys 38, 215 (1983). URL http://stacks.iop.org/0036-0279/38/i=4/a=R11.

[71] Kessler, D. A., Ner, Z. & Sander, L. M. Front propagation: precursors, cutoffs, and structural stability. Physical Review E 58, 107 (1998).

[72] Mikhailov, A. S., Schimansky-Geier, L. & Ebeling, W. Stochastic motion of the propagating front in bistable media. Physics Letters A 96, 453–456 (1983).

[73] Rocco, A., Casademunt, J., Ebert, U. & van Saarloos, W. Diffusion coefficient of propagating fronts with multiplicative noise. Physical Review E 65, 012102 (2001).

[74] Meerson, B., Sasorov, P. V. & Kaplan, Y. Velocity fluctuations of population fronts propagating into metastable states. Physical Review E 84, 011147 (2011).

[75] Panja, D. Effects of fluctuations on propagating fronts. Physics Reports 393, 87–174 (2004).

[76] Reguera, D., Reimann, P., Hänggi, P. & Rubi, J. Interplay of frequency-synchronization with noise: Current resonances, giant diffusion and diffusion-crests. EPL (Europhysics Letters) 57, 644 (2002).

[77] Matthysen, E. Density-dependent dispersal in birds and mammals. Ecography 28, 403–416 (2005).

[78] Liu, C. et al. Sequential establishment of stripe patterns in an expanding cell population. Science 334, 238–241 (2011).

[79] L’heureux, N., Lucherini, M., Festa-Bianchet, M. & Jorgenson, J. T. Density-dependent mother-yearling association in bighorn sheep. Animal Behaviour 49, 901–910 (1995).

[80] van Ditmarsch, D. et al. Convergent evolution of hyperswarming leads to impaired biofilm formation in pathogenic bacteria. Cell reports 4, 697–708 (2013).

[81] Phillips, B. L., Brown, G. P., Webb, J. K. & Shine, R. Invasion and the evolution of speed in toads. Nature 439, 803–803 (2006).

[82] Phillips, B. L. The evolution of growth rates on an expanding range edge. Biology Letters 5, 802–804 (2009). URL http://rsbl.royalsocietypublishing.org/content/5/6/802. http://rsbl.royalsocietypublishing.org/content/5/6/802.full.pdf.

[83] Korolev, K. S. et al. Selective sweeps in growing microbial colonies. Physical biology 9, 026008 (2012).

[84] Wolf, D. E. Wulff construction and anisotropic surface properties of two-dimensional eden clusters. Journal of Physics A: Mathematical and General 20, 1251 (1987). URL http://stacks.iop.org/0305-4470/20/i=5/a=033.

[85] Savit, R. & Ziff, R. Morphology of a class of kinetic growth models. Phys. Rev. Lett. 55, 2515–2518 (1985).

[86] Nahum, A., Vijay, S. & Haah, J. Operator spreading in random unitary circuits. Phys. Rev. X 8, 021014 (2018).

[87] Fisman, D. N. Seasonality of infectious diseases. Annu. Rev. Public Health 28, 127–143 (2007).

[88] Johnson, D. M., Liebhold, A. M., Tobin, P. C. & Bjornstad, O. N. Allee effects and pulsed invasion by the gypsy moth. Nature 444, 361–363 (2006).

[89] Sullivan, L. L., Li, B., Miller, T. E. X., Neubert, M. G. & Shaw, A. K. Density dependence in demography and dispersal generates fluctuating invasion speeds. Proceedings of the National Academy of Sciences (2017).

[90] Weinberger, H. F. On spreading speeds and traveling waves for growth and migration models in a periodic habitat. Journal of mathematical biology 45, 511–548 (2002).

[91] Deforet, M., Carmona-Fontaine, C., Korolev, K. S. & Xavier, J. B. A simple rule for the evolution of fast dispersal at the edge of expanding populations. arXiv preprint arXiv:1711.07955 (2017).

[92] Marquet, C., Peschanski, R. & Soyez, G. Consequences of strong fluctuations on high-energy QCD evolution. Phys. Rev. D 73, 114005 (2006).

[93] Douglas, J. F., Efimenko, K., Fischer, D. A., Phelan, F. R. & Genzer, J. Propagating waves of self-assembly in organosilane monolayers. Proceedings of the National Academy of Sciences of the United States of America 104, 10324–10329 (2007).

[94] Ben-Jacob, E., Cohen, I. & Levine, H. Cooperative self-organization of microorganisms. Advances in Physics 49, 395–554 (2000).

[95] Gerlee, P. & Nelander, S. The impact of phenotypic switching on glioblastoma growth and invasion. PLoS Computational Biology 8, e1002556 (2012).

[96] Lavrentovich, M. O., Korolev, K. S. & Nelson, D. R. Radial domany-kinzel models with mutation and selection. Physical Review E 87, 012103 (2013).

[97] Rouzine, I. M., Wakeley, J. & Coffin, J. M. The solitary wave of asexual evolution. Proceedings of the National Academy of Sciences 100, 587–592 (2003).

[98] Aronson, D. G. & Weinberger, H. G. Nonlinear diffusion in population genetics, combustion and nerve propagation Lectures Notes Math, vol. 446 (Springer, New York, 1975).

[99] Bender, C. M. & Orszag, S. A. Advanced Mathematical Methods for Scientists and Engineers I (Springer Science & Business Media, 1999).

